# Genomic Footprints of Bottlenecks, Isolation, and Inbreeding: A Case Study of Two Vulture Cohorts in India

**DOI:** 10.64898/2026.04.30.721611

**Authors:** Manas Shukla, Dau Lal Bohra, Babu Rao, Laxmi Narayan, Shashi Kiran, Vivek Thakur

**Author notes:** **Author emails** Manas Shukla Dr Shashi Kiran Dr Daulal Bohra Dr Babu Rao Dr Laxmi Narayan. **indicates Corresponding author:** Dr Vivek Thakur.

## Abstract

Genomic erosion as a manifestation of small effective population size (N_e_) and consanguinity subverts long-term perpetuation of threatened species by compromising their adaptive potential; however, the integration of genomics remains limited in applied conservation efforts to guide priorities. This study combines non-invasive sampling, double-digest Restriction site-associated DNA sequencing (ddRAD), and population-genomic analyses to assess genetic health in two vulture assemblages-mixed wild enclosure and captive breeding cohorts. Both the geographical locations exhibit signs of populations in distress: low genetic diversity and abundant intermediate-length ‘runs of homozygosity’ (RoH), consistent with long-term reduced N_e_ plus recent demographic isolation. Our demographic model runs favoured ancient migration (AM) topology characterised by an ephemeral window of gene flow, taken over by a prolonged population separation period. The mutation quantification results from approximately 59,000 outgroup-polarised SNPs reveal higher additive burden and more homozygous-derived sites in BKN. However, this was later traced to low-impact and non-coding variants rather than a surge in the loss-of-function (LoF) alleles. The data support a genomic profile that carries an elevated risk from polygenic/aggregate deleterious burden in BKN despite a scarcity of high-impact mutations. By highlighting the disconnect between genetic resilience and demographic recovery, our results accentuate the need to incorporate genomics-informed inbreeding and monitoring programs, while also focusing on reducing anthropogenic mortality with genetic augmentation.

## Introduction

Nearly 15% of the global extant birds are threatened with extinction, and recent decades have witnessed precipitous population decline in an alarmingly large number of species (Li 2014; Storch et al. 2023). The incessant avian decline has been majorly attributed to habitat degradation, climate change, pesticide abuse, and hunting (Li 2014). Among these, the vulture population collapse observed in the late 20^th^ century was one of the most remarkable declines of the time, chronicled by over 99% population losses in three *Gyps* residents of India (Adawaren et al. 2024; Swan 2006). This outrageous plunge was later ascribed to uric acid accumulation in the blood and subsequently, visceral gout due to toxic levels of diclofenac (an NSAID) in the blood (Lierz 2003; Oaks 2004; Swan 2006). A statutory ban on veterinary use of diclofenac followed in 2006, but the implementation has been poor, and diclofenac still poses a threat to vultures (Ghosh 2024).

Vultures, being obligate scavengers, are pivotal to ecosystem sanitation and have profound, measurable impacts on human health and livelihoods by rapidly removing the carcasses that would otherwise sustain pathogen reservoirs. Their collapse was followed by an increased carrion persistence associated with a documented rise in feral dog bites and rabies spread in humans, with estimates attributing hundreds of thousands of excess deaths and morbidity events (Biswas 2025; Frank and Sudarshan 2024). These ecological and societal feedbacks underscore that vulture conservation must be both a biodiversity and a human well-being priority.

Demographic recovery after such crashes is necessary but does not entail genetic recovery, further supported by the theory of drift debt in small populations (Dehasque 2024; Gilroy et al. 2017; Ochoa et al. 2020). Empirical studies in the past emphasize that small effective population sizes (N_e_) inadvertently reduce mate choices, leading to increased inbreeding due to the *Allee* effect, further depleting genetic variation and weakening selection’s ability to purge mildly deleterious variants even after census size rebounds (Drake and Kramer 2011; Simons et al. 2014). The literature indicates that such purging can occur, but recovery of standing genetic variation may require many generations and, for many species, may be effectively irreversible on management timescales (Bemmels 2025; Dehasque 2024; Star and Spencer 2013). Hence, demographic trends alone overestimate recovery, indicating conservation assessments must incorporate genomic baselines that directly measure genetic erosion and functional burden.

Genomic monitoring is therefore an urgent need for vulture conservation. While whole-genome sequencing remains the gold standard for comprehensive genetic analyses, alternate approaches also provide a cheaper, scalable alternative when broad genome representation is captured across many independent loci (Luca et al. 2011; Theissinger 2023).

Here, we evaluate whether threatened Indian vultures exhibit signatures of long-term genetic erosion that could compromise recovery, and we present an evidence-driven framework for routine genetic monitoring. The Indian subcontinent houses nine vulture species, five of which remain endangered (EN) or critically endangered (CR) to date (***Supplementary Table 1***) ([3]). Interestingly, the endangered species are all residents, while the species with the least-concerned (LC) tag are either migratory or partially migratory vultures.

This study investigates the ***hypothesis*** that the endangered Indian vultures face preordained genomic erosion as a consequence of prolonged demographic decline. The study followed an integrative framework involving non-invasive feather sampling involving representatives of both endangered/critically endangered (EN/CN) and least-concerned (LC), followed by population genomic analysis. The framework allowed us to: (1) reliably capture the population-genetic metrics relevant to conservation, (2) quantify per-sample mutation burden and recessive load, and (3) interpret load differences in light of demographic history, inbreeding, and conservation. By combining inferences from all the conservation-relevant metrics, this framework intends to provide robust, actionable genomic baselines for vulture conservation and to demonstrate a scalable framework for monitoring other threatened avian taxa.

## Methodology

### Sampling and Ethics Statement

Feathers were collected opportunistically as moulted materials from the Vulture Conservation Centres and/or Breeding Centres of India over a three-month sampling window (December 2024 to February 2025). The “Jorbeer Vulture Conservation Centre”, Bikaner, Rajasthan (hereafter “BKN”), and “Vulture Breeding Centre” (VBC), Nehru Zoological Park, Hyderabad, Telangana (hereafter “HYD”) constituted the actual sampling locations (***Supplementary Fig. S1***). All sampling was conducted under the relevant institutional and state permits (see ***Supplementary Fig. S2***).

Since NZP only harbours White-rumped Vultures (*Gyps bengalensis*), all HYD feathers were considered as originating from *G. bengalensis*. While the BKN, being an open enclosure, supports multiple vulture species, feathers from Egyptian Vulture (*Neophron percnopterus*) and Eurasian griffon Vulture (*Gyps fulvus*) were collected preferentially, although field identification of species and individual assignment could not be confirmed with certainty from feather morphology alone. Therefore, all BKN samples were treated as a spatial sample set throughout the study in an attempt to maintain transparency and to avoid circular species assignment.

### Genomic DNA (gDNA) Extraction

We compared two DNA extraction protocols established for feathers: a Phenol-Chloroform-Isoamyl Alcohol (PCI)-based protocol for calamus tips and a silica-column kit-based protocol for the dry clot appearing outside the superior umbilicus region (Bantock et al. 2008; Brett et al. 2008; Horváth et al. 2005; Margeta et al. 2021; Presti et al. 2013; Volo et al. 2008) (see ***Supplementary Fig. S3***). Given its poor success rate, we adopted the **clot-based** protocol for DNA extraction (Aksel and Arslan 2023; Horváth et al. 2005; Ozdemir et al. 2024; Presti et al. 2013). A breakdown of DNA quantity and purity can be inferred from the ***Supplementary Table S3***. Since the feathers were collected opportunistically, there remained an ambiguity over their moulting period, likely a factor affecting PCI-based DNA extraction. DNA from the clot was extracted with Nucleospin® Tissue DNA kits (Lot: 2502-005 and 2501-002; Germany), assimilating modifications discussed by Dr Presti’s group (Presti et al. 2013). A nanodrop spectrophotometer was used to assess the quantity of the extracted DNA from the kits. The DNA was further evaluated for its integrity by using Agarose gel electrophoresis (1% gel), Lonza, Belgium, and visualized in the Vilber Bio-Print Documentation system. See *supplementary methods* for wet and dry-lab QC.

### Genotyping

DNA sequencing libraries were prepared by *Nucleome Informatics Pvt. Ltd*. (Telangana, India) using a restriction digestion protocol, utilizing the Kapa Hyperplus kit (Manufacturer: *Roche*). The restriction enzymes MseI (frequent cutter) and NlaIII (rare cutter) were used to allow fragments in the window 300–500 bp. An Agilent 2100 Bio-analyser was used for library length assessment. Each library was constructed from one feather DNA extract, where unique libraries contained individual barcodes, and sequencing was performed using Illumina’s NovaSeq 6000 S4 reagent kit V1.5 (2×150bp, PE).

### Dataset Curation and Workflow Finalisation

See *supplementary methods*.

### Heterozygosity and Inbreeding Assessment

To gauge genetic diversity within each demographic cluster, we calculated standard population genetic summary statistics from filtered SNP datasets. Only biallelic SNPs were considered, while sites with missing genotypes or monomorphic loci were excluded from the analysis (Natesh 2017). Individual-level heterozygosity was estimated using the -het function in PLINK v2.00a3 (Chang et al. 2015). The full SNP dataset, comprising 86,140 high-quality biallelic SNPs, was used for these analyses to preserve the maximum amount of genome-wide information. Variants and genotypes passing the quality threshold criteria (QUAL ≥ 30, DP ≥ 10, GQ ≥ 20, MQ ≥ 40, MAF ≥ 0.03, and missingness ≤ 0.7) were retained, and all the individuals were treated as founders.

Runs of Homozygosity (RoH) were estimated within individuals across all the population clusters by availing PLINK v1.90b6.24 (Purcell 2007) with a window-based approach to detect extended tracts of homozygosity consistent with autozygosity (Ceballos et al. 2018b; McQuillan 2008). The analysis was performed using the filtered binary genotype files (.bed/.bim/.fam) for each cluster. We enabled the –*homozyg* function in PLINK where RoHs were identified using sliding-window approach, requiring a minimum RoH length of 1 Mb containing at least 50 homozygous SNPs, with a maximum inter-SNP gap of 1 Mb and a minimum SNP density of one SNP per 50 kb; up to one heterozygous genotype and five missing genotypes were permitted per window, and RoH were called only when windows contained at least 50 SNPs.

We adopted a different set of parameter values more tailored for analysing low-coverage data, predicated on the recommendations from past studies (Berghöfer et al. 2021; Ceballos et al. 2018a; Duntsch et al. 2021; Lavanchy and Goudet 2023). The individual RoH segments were binned into size categories mirroring their likely age: very small (500-1000 kb), small (1000-2000 kb), medium (2000-5000 kb), and long (>5000kb).

### ADMIXTURE and Principal Component Analysis (PCA)

Model-based genetic ancestry estimates were performed using ADMIXTURE (Alexander et al. 2009) on both the LD-pruned SNP sets (both pruned_relaxed and pruned_stringent PLINK *bfiles* produced during the dataset curation run; ***see Fig. 1***). Each pruned file was copied to an integer-mapped version before analysis to overwrite the NCBI chromosome codes. ADMIXTURE was run for K = 1-6 for each pruned file with 5 independent replicates per K, using different random seeds and ten-fold cross-validation (--CV = 10) to assess model stability and select the best run with the least CV error; replicate P/Q metrics were inspected for convergence (quasi-newton algorithm) and consistency across runs. PLINK was used for LD-pruning and file format handling.

**Fig 1.**
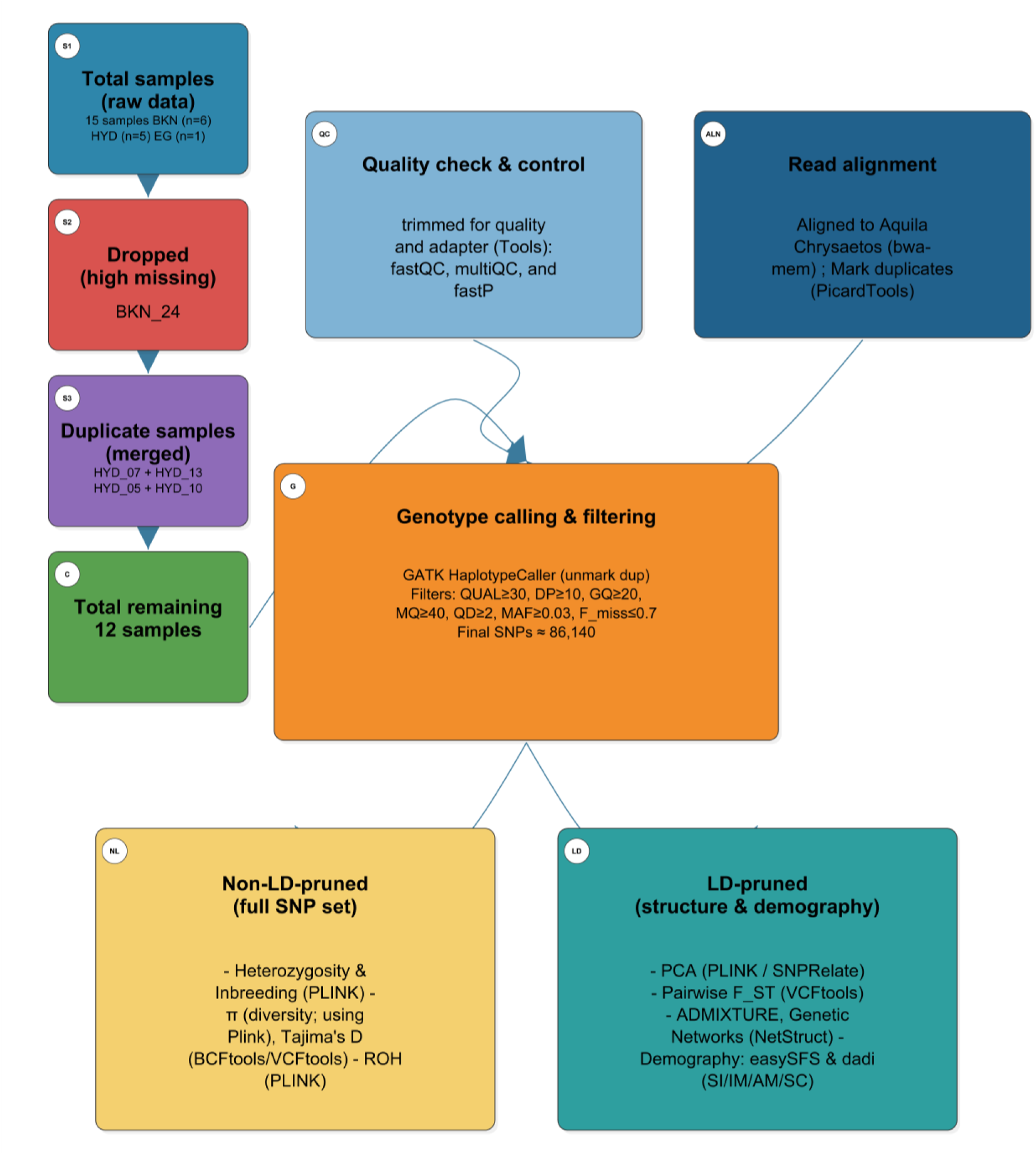
Final analysis workflow and filtering summary. Project workflow from raw samples through QC, alignment, and genotype-calling (GATK HaplotypeCaller), with downstream branches for LD and non-LD pruned SNP sets. Key genotype filters are listed in the ‘genotype calling and filtering’ section.

Principal Component Analysis (PCA) was computed from the same LD-pruned VCFs using the R package SNPRelate: VCFs were converted to GDS (snpgdsVCF2gds), and snpgdsPCA was used to derive eigenvectors and eigenvalues (Patterson et al. 2006; Zheng et al. 2012a). Sample population mapping was applied, and PCA scores were saved and plotted (PC1 vs PC2 and PC1 vs PC3) with population colour mapping for visualisation and outlier identification. Variance was incorporated for the PCs from eigenvalues to support the interpretation of clusters and population structure.

### Population Differentiation Analysis (Pairwise F_ST_ Measure)

To evaluate genetic differentiation among the population clusters, pairwise FST analyses were performed on both the LD-pruned SNP sets using a custom pipeline integrating bcftools v1.21 and VCFtools v0.1.16. Sample lists (*pop* files) were extracted from *nosex* files generated by PLINK v1.90b6.24, followed by merging and indexing the normalised VCF files across all the clusters for the LD-pruned sets. Population pairwise FST (Weir and Cockerham estimator) was calculated using the –weir-fst-pop option on pruned VCFs and per-population sample lists; both the mean and weighted FST values recorded by vcftools were saved for summaries and plotting (Knaus and Grünwald 2017; Paradis et al. 2004; Weir and Cockerham 1984).

Individual-level pairwise F_ST_ was gauged in R from the pruned VCF genotypes (GT→numeric 0/1/2) using a per-site genotype-based estimator and summarised as mean per-sample F_ST_ across sites (a minimum overlapping site filter was enforced; default min_sites=100). The script constructs an individual X individual F_ST_ matrix, heatmaps, and MDS/PCoA from the pairwise matrix, besides generating per-sample summary tables.

### Genetic Similarity Networks

Genetic network inferences were built on LD-pruned genotype matrices (PLINK→SNPRelate) by calculating pairwise distances as the mean absolute genotype difference (|g_i_ – g_j_|/2) across sites and converting this to a similarity matrix *S* = **1-*D*** (Clauset et al. 2004; Dyer and Nason 2004; Zheng et al. 2012b). Networks were constructed for a sweep of genetic similarity thresholds (edge present if S ≥ threshold), with replicates used to assess network and threshold robustness for an effective identification of stable sample groupings. Communities were uncovered via a fast greedy modularity algorithm (*igraph*), and per-threshold summaries (node, edges, degrees, modularity), community membership, and per-sample metrics (mean distance, nearest neighbour) were recorded while figures were subsequently rendered with weighted edges (Csárdi and Nepusz 2006).

### Nucleotide Diversity Estimation

To appraise the patterns of genetic diversity, we estimated nucleotide diversity (π) per population using VCFtools (v0.1.16) –windows-pi and –site-pi on the full, non-LD pruned SNP dataset (Danecek 2011) produced by our curation pipeline (***Fig. 1***). In an effort to obtain stable windowed π estimates for a shallow ddRAD dataset, window sizes were dynamically selected from observed SNP density such that each window contained approximately 10 SNPs (Zhang et al. 2023). We used a custom Python script for our calculations, avg-bp-per SNP was computed from the VCF (or a fallback value of 14,322 bp when contig lengths were unavailable), and the window length (bp) was computed as round (avg_bp_per_snp × target_snps).

Windows were limited to a minimum of 5 Kb to a maximum of 5 Mb, and the step size was chosen to be half the window length to allow partially overlapping windows. This approach balances noise from small windows against spatial resolution in reduced-representation data. Population sample lists (*keep* files) were fed to vcftools to evaluate both windowed π and per-site π for each population.

The resultant windowed **π** files (per-population) were merged into a single table for comparative plotting and downstream analysis. Basic summary statistics (no. of windows, mean **π**, median **π**, standard deviation of **π**) were computed from the windowed output for each population, and per-site **π** files were retained for locus-level inspection. An R plotting script was run to visualise windowed **π** across populations and produce relevant figures.

#### Tajima’s D Estimation

We next investigated the potential deviations from neutral evolution, inferring demographic history or selective sweep signatures, by computing Tajima’s D value across genomic windows using VCFtools v0.1.16.

We employed two complementary approaches to evaluate Tajima’s D from the final filtered VCF (86,140 SNPs): a) non-overlapping windows computed directly with vcftools (--Tajima D) for the full dataset, and b) sliding (50% overlap) windows generated with bedtools (Quinlan and Hall 2010) and evaluated per-window by extricating the regions with bcftools and running vcftools **– *TajimaD*** on each region. To optimise resolution and statistical stability for ddRAD data, windows were tailored to match the SNP density data (default 1,432,000 bp ≈ 100 SNPs/window; step = 716,000 bp). Further windows with no variants were recorded with n_variants = 0 and TajimaD=NA, and vcftools nan outputs were converted to NA in downstream CSVs, which were later used in R v4.4.1 to visualise the distribution. Window length and population sample decisions from the pilot run were applied before computation.

### Mutation Load Quantification

We annotated variants and estimated mutation load by developing a custom SnpEff database for *Aquila chrysaetos* from our project reference FASTA and GTF, extracting CDS and protein sequences with *gffread* and running the SnpEff JAR (v5.3a) to build the genome and annotate the filtered VCF. Further, for each variant, the SnpEff ANN field was parsed to assign a **worst predicted impact (HIGH, MODERATE, LOW, MODIFIER) and consequence term (e.g**., **stop_gained, splice_acceptor, missense_variant)**. Per-sample genotypes were converted to alternate-allele dosages (0 for 0/0, 1 for heterozygotes, 2 for homozygous alt) with missing genotypes excluded, and homozygous-derived counts were noted separately. ***See Supplementary Methods*** for steps taken to minimize interspecific and related biases.

We estimated uncertainty by site-level bootstrap (resampling sites; default 2,000 replicates) to produce 95% CIs for means and per-impact homozygous counts. Between-group tests employed non-parametric tests (two-sided Wilcoxon rank-sum) with medians and median differences reported as effect sizes, while enrichment in sharing categories was assessed by permutation tests (10,000 permutations) with Benjamini-Hochberg FDR correction.

### Demographic Modelling and Inferences

Demographic modelling was performed from a folded joint site-frequency spectrum (SFS) computed with a pipeline integrating ANGSD and realSFS with ∂a∂i demographic model for fitting (Gutenkunst et al. 2009; Korneliussen et al. 2014). Concisely, BAMs were reorganised into population-specific lists, and ANGSD (samtools GL model, GL -1) was directed with conservative base/mapping filters (e.g., MinMapQ=30, minQ=20, only_proper_pairs, BAQ) and population-specific -minInd thresholds to produce saf.idx files. realSFS was used to compute 1D SFS for each population and the folded 2D joint SFS (realSFS fold), exported as a matrix of numeric counts and converted to ∂a∂i input (rows = 2nBKN+1, cols = 2nHYD+1).

Model fitting was performed via a custom Python script implementing a resume capable ∂a∂i pipeline that a)-reads the folded-joint SFS, b)-examines multiple two-population models, namely isolation with continuous migration (IM), secondary contact (SC), strict isolation (SI), and ancient migration (AM), and c)-uses a two-stage optimisation strategy (coarse multi-start searches followed by refined optimisation) with extrapolation grids to stabilise likelihoods. Fitted model SFS were scaled by optimal theta and assessed by multinomial log-likelihood; parameter uncertainty was scanned by parametric (multinomial) bootstraps (-nboot=100), yielding bootstrap parameter distribution and 95% confidence intervals. Models were compared using Akaike Information Criterion (AIC) and *Akaike* weights; diagnostics (*observed vs expected SFS, residuals, parameter violin/histogram plots, pairwise scatter, and bootstrap correlation heatmap*) were recorded alongside fit summaries.

## Results

### Dry Clots in Tail Feathers Yielded Superior DNA but had Poor Integrity

Freshly moulted feathers were sampled from two of the vulture breeding and conservation sites within India (***Supplementary Fig. S1, Table S1)***, differing in terms of species composition and enclosure settings (open/captive). A minimum of 6 individuals per site (BKN/HYD) was adhered to for useful inferences about the population genetic metrics (refer to ***Supplementary Table S2*** for the extraction workflow and per-species information).

We found significant improvement in gDNA extraction when only rectrices (tail feathers) were considered (42.96 ± 20.40 ng/µL; n = 18) over alternate approaches, across all vulture species tested. This observation is consistent with the previous findings (Horváth et al. 2005; Volo et al. 2008), which also reported superior DNA yield from tail feathers. Based on the quality and quantity of extracted genomic DNA, 16 samples were selected for further validation, library preparation, and genotyping (***Supplementary Table S3***).

The ddRADseq genotyping generated sequence data with a mean of ≈ **590.64 MB** bases (*Median* ≈ *591.32 MB; Min*. ≈ ***394 MB***; *Max*. ≈ ***785 MB***) and moderate variability among the samples (Coefficient of Variation, CV = O’*/mean = 103.46 MB/*590.64 MB = **0.16**) (***Supplementary Table S6)***.

### Pilot-run-based Curation Addressed the Grouping or Genotyping Issues

Based on the pilot run (see *Supplementary Methods*), a few issues were observed which were addressed by curation: a sample with high missingness was dropped (BKN_24), the one failing to form a group was considered an outlier (BKN_35 aka BKN_UV_o_01), and duplicate samples were merged (N=2).

After curation, joint genotyping identified common and unique variants in the remaining 12-sample dataset (see the workflow in ***Fig. 1***). Three analysis sets were obtained from the final VCF file: **a non-LD pruned SNP set** of 86,140 SNPs, and **two LD-pruned sets** (one *relaxed set* containing 9,241 SNPs and another *stringent set* of 4,042 SNPs) for population structure and demography-based inferences (refer to ***Supplementary Fig. S10*** for comparisons before and after pruning). See *Supplementary Results* for more information.

### ROH Size Spectrum Reveals Overall Higher Recent Autozygosity in BKN Samples

The assumption that extreme bottlenecks and low long-standing population effective size (N_e_) entail ineluctable negative consequences of inbreeding in the affected populations formed the basis of this analysis. Of the two approaches for inbreeding estimates, the PLINK-based approach produced very low observed heterozygosity values and very large inbreeding coefficients (**F** ≈ **0.89–0.91** for most samples; see ***Fig. 2a***). These **F** estimates clearly indicate a large depletion of heterozygotes relative to Hardy-Weinberg expectations on the assayed marker set.

**Fig 2.**
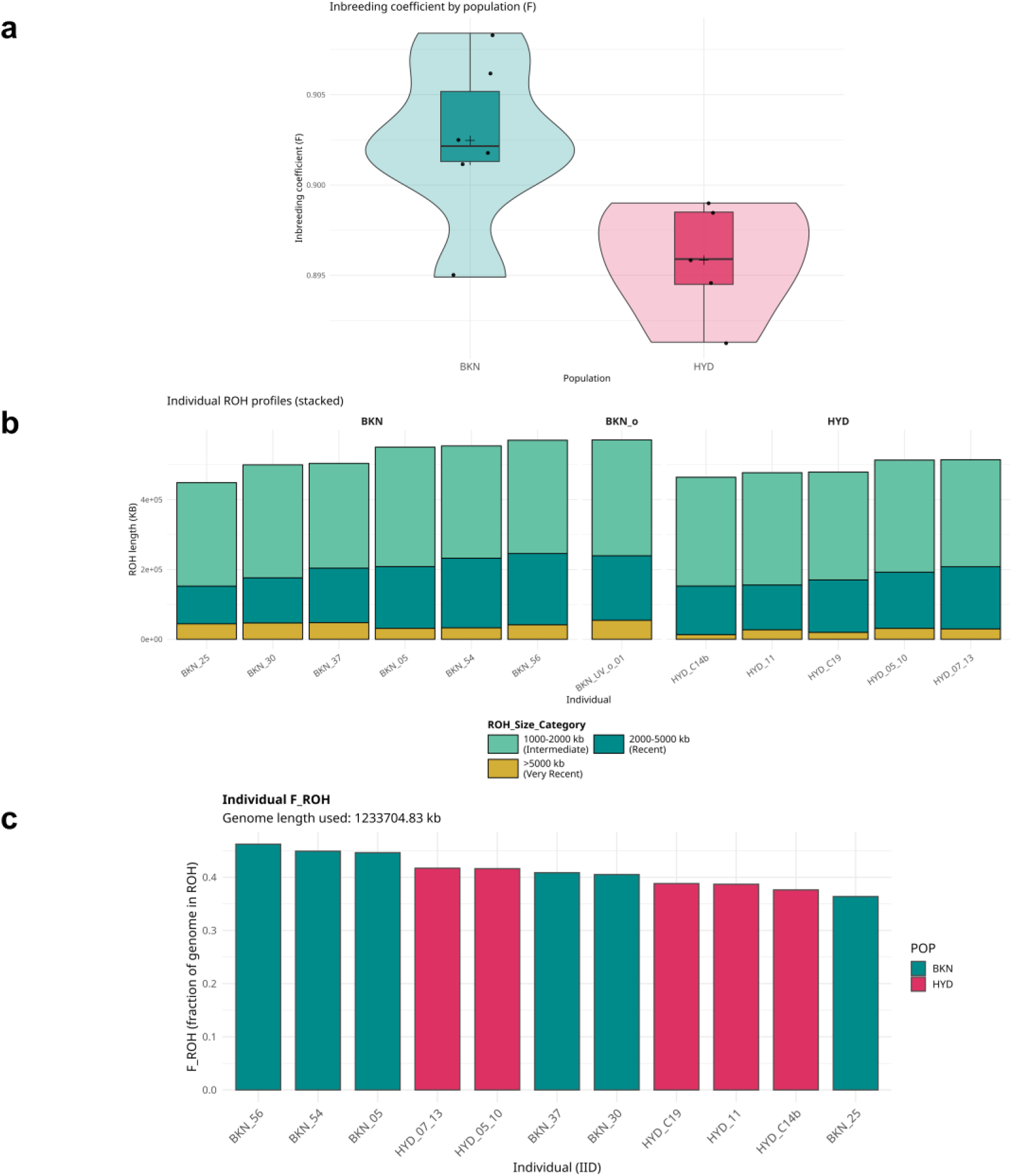
Inbreeding Estimation. (a) Inbreeding coefficient *F* by population. Violin plots comparing per-sample inbreeding coefficients between populations; BKN represents a higher median F and a narrower distribution than HYD, indicating greater genome-wide homozygosity. The white diamond shows the mean value for the distribution. **(b) Individual ROH profiles:** Stacked bar-plot reflecting the proportion of homozygous tracts derived as a consequence of historical/recent mating among relatives **(c) F**_**ROH**_ **estimation:** Bars show the genomic inbreeding coefficient value estimated as a fraction of total homozygous tract length to the callable genome length.

We investigated recent and historical autozygosity by calling runs of homozygosity (ROH) and summarising ROH by size classes, which also reflect the relative timing of the inbreeding events (1-2 Mb = older autozygosity; 2-5Mb = recent autozygosity; >5Mb = very recent autozygosity; for more information, refer to ***Supplementary table S7***). Summed across ROH bins, the mean total ROH per individual was slightly higher in **BKN** than in **HYD** (BKN ≈ 521 Mb per individual; HYD ≈ 490 Mb per individual). The outlier from the BKN group had a total ROH (≈571 Mb) at the top of the distribution and should be interpreted as a single-sample observation. The population level breakdown of the ROH classes can be inferred from ***Fig. 2a*** (also see ***Supplementary Fig. S11a-b)***. *See supplementary results for a discussion on the methodological reasons for quantitative differences in the two approaches of heterozygosity estimation*.

The large number of intermediate-length ROH segments per individual signifies that a considerable fraction of autozygosity derives from *older* demographic processes (persistently small long-term N_e_ or historical bottlenecks), while the presence of long tracts is a diagnostic of *recent* related mating or very recent severe reductions in local N_e_ (Ceballos et al. 2018b; McQuillan 2008b). The relatively larger burden of *recent* ROH in BKN thus indicates that **BKN has experienced more recent inbreeding or recent demographic isolation than HYD**.

To infer genomic inbreeding, we next computed F_*ROH*_ values (see ***Fig. 2c;*** also refer to ***Supplementary Fig. S11c***). **HYD** samples had a **mean F**_***ROH***_ **of 0.397 (± 0.0185)**, whereas **BKN** samples (excluding BKN_UV_o_01) boasted even higher coefficients (**mean F**_***ROH***_ ≈ **0.422 ± 0.0369**). The values reported here imply that a large proportion of the callable genome is in homozygous runs, reiterating severe long-term reductions in the effective population size and prolonged population isolation of the sampled groups.

### Species Composition in Different Enclosure Settings (Open/Captive) Drive Observed Genetic Clustering

To establish whether geographical mapping coincides with genetic grouping, we examined population structure across 12 samples using four approaches: principal component analysis (PCA), pairwise and group F_ST_ estimates, model-based clustering (ADMIXTURE), and genetic-similarity networks. All analyses used LD-pruned SNPs (see Methods). PCA, ADMIXTURE, and the network results are concordant in revealing two main clusters that map to their geographical sampling locations (BKN: Captive Assemblage; HYD: Open Enclosure), with the outlier sample acting as a distinct entity or third cluster depending on the method.

PCA of the pruned samples separates them by sampling site primarily by **PC1** (**39.9% of variance**; see ***Fig. 3a***). Additionally, pairwise F_ST_ between all individuals was computed (Weir & Cockerham estimators were used -- see *Methods*). Individual pairwise F_ST_ values within BKN are extremely low (most values of the order ≈ 10^−6^ to 10^−4^), demonstrating high similarity among the group members. In comparison, the pairwise F_ST_ for intergroup pairs is larger (∼0.005-0.007 for many pairs), reflecting measurable differentiation at our SNP set. The outlier displays low pairwise F_ST_ with most BKN individuals (≈ 0.0003-0.0009) and slightly higher values to HYD individuals (≈ 0.0027-0.0031), further reinforcing the findings from PCA (see ***Fig. 3b***).

**Fig 3.**
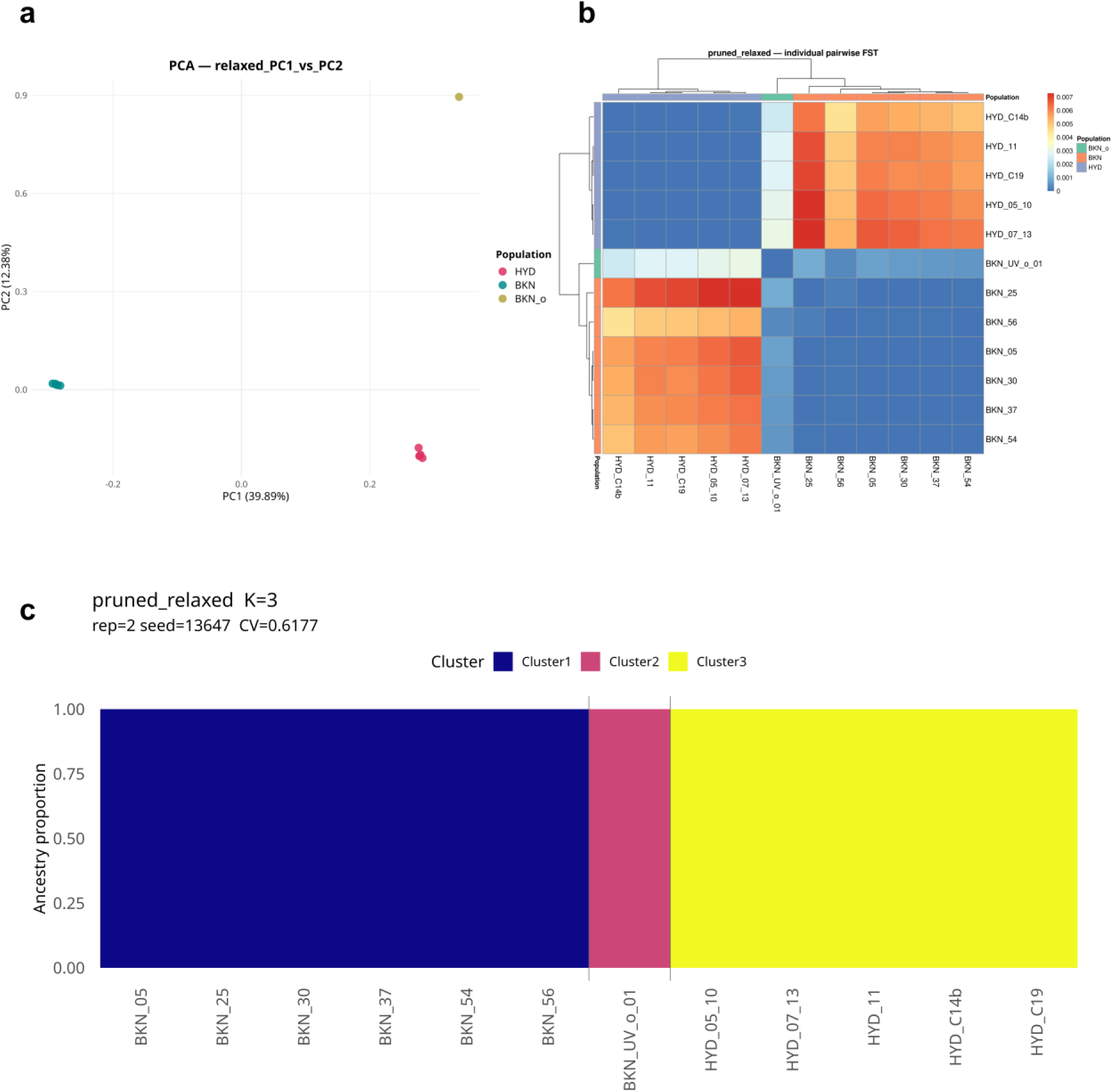
Population Differentiation/Divergence. (a) Principal Component Analysis (PC1 vs PC2). Multi-dimensional scaling of 12 samples (LD-pruned ‘relaxed’ SNP-set) showing PC1 (39.9% variance) and PC2 (12.4%). Samples cluster by geographical locations (BKN, HYD) with one BKN sample (BKN_UV_o_01) appearing as an outlier. **(b) Individual pairwise F**_**ST**_ **heatmap** (Weir and Cockerham estimator) illustrates very low within-BKN differentiation (≈10^−6^-10^−4^) and larger inter-group values (≈0.005-0.007). **(c) ADMIXTURE bar-plot** (K=3, best supported by CV; K=1-6 tested) reiterates two major ancestry components corresponding to sampling sites and a distinct signal associated with BKN_UV_o_01.

The next metric of population divergence was provided by model-based clustering using the tool ADMIXTURE (see Methods). ADMIXTURE was run for K = 1-6 (six replicates each; for breakdown of each replicate run, see ***Supplementary Table S8***). Cross-validation (CV) and replicate consistency (standard deviation in CV) establish K = 3 as the best balance of fit and stability in the pruned dataset, further corroborating the findings of PCA (see ***Fig. 3c;*** also refer to ***Supplementary Fig. S13*** for details on all the K runs). At K = 3, the major pattern is contained in three group structures: a BKN cluster, a HYD cluster, and a singleton group constituted by *BKN_UV_o_01* (it forms either a distinct cluster or shows mixed ancestry depending on K and replicate, ‘sticking out’ in all runs). Because BKN samples contained one or two species of vultures (likely Egyptian vultures and/or Eurasian griffon vultures) while HYD consisted of White-rumped vultures solely, part of the signal may indicate species-level differentiation. The outlier may represent a different species/lineage or an individual with partial ancestry from the HYD cluster.

In another attempt to capture divergence among the samples, we constructed weighted genetic-similarity networks from the similarity matrix (S = 1 – D, D = pairwise genotype distance), and explored a dense set of thresholds (t). At low threshold (**t = 0.10**), the **network is dense** (66 edges & 12 nodes) and segregates into 2 communities with low modularity (0.021; low sub-group confidence). As weak edges are removed, modularity increases: t = 0.70 yields 30 edges and modularity ≈ 0.5, t = 0.87 yields 14 edges and 3 communities (modularity ≈ 0.480), and t = 0.90 yields 14 edges and 4 communities (modularity ≈ 0.490; see ***Fig. 4*** and ***Supplementary Fig. S14***). Even at higher thresholds, samples within sites remai this indicates high within-site similarity. The behaviour of the outlier sample across thresholds emulates the ADMIXTURE/PCA findings: it can be assigned to a distinct community or a group with HYD, depending on the threshold.

**Fig 4.**
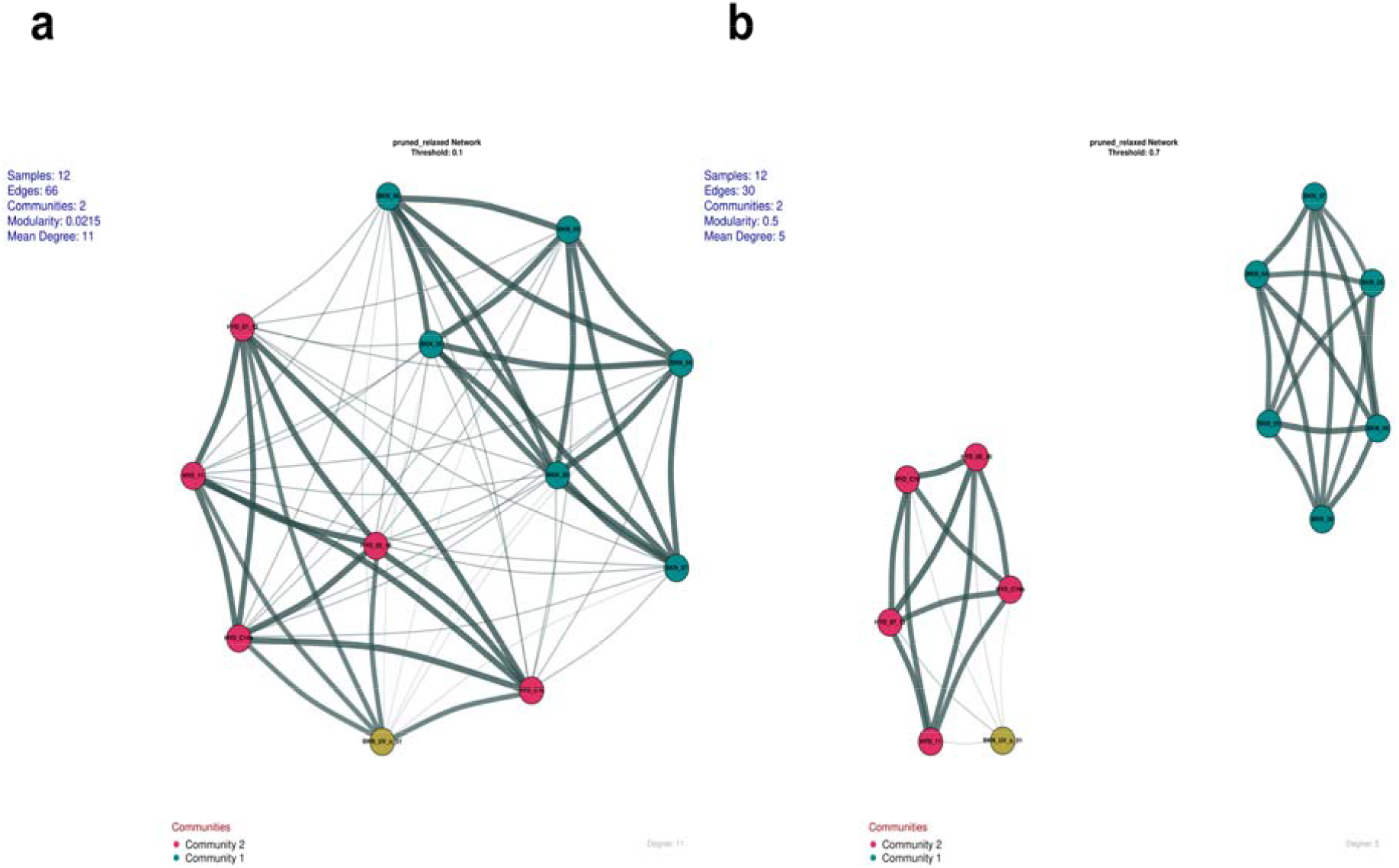
Weighted genetic similarity networks. Community detection algorithm employed on the basis of genetic similarity (S = 1 – D, where D = pairwise genotype distance), visualized across edge-thresholds t=0.10 and t=0.7. At a low threshold, the network is dense and poorly modular; as weak edges are expelled, modularity increases, and within-site connectivity remains high.

### Low Nucleotide Diversity Consistent with Recent Bottlenecks, particularly in BKN

To quantify within-population genetic diversity and test for departures from neutral evolutionary expectations, thereby revealing the demographic history and selective pressures shaping the population, we estimated nucleotide diversity (**π**) and Tajima’s D in optimized windows (see Methods). The **mean** windowed **π in the BKN group** was ***5.42 × 10***^***-6***^ (median = 3.70 × 10^−6^; n = 8,813 windows) **and *6.05 × 10***^***-6***^ in HYD samples (median = 3.88 × 10^−6^; n = 8,408 windows**) (*Fig. 5a*)**. Mean Tajima’s D was estimated to be **−0.51 in BKN** (median −0.54, SD = 0.68, n = 1,687 windows) and **-0.20 in HYD** (median −0.21, SD = 0.64, n = 1,599 windows) (***Fig. 5b***). The results together reveal consistent, species- and site-level hallmarks of reduced diversity, skewed allele frequencies and recent demographic perturbation.

**Fig 5.**
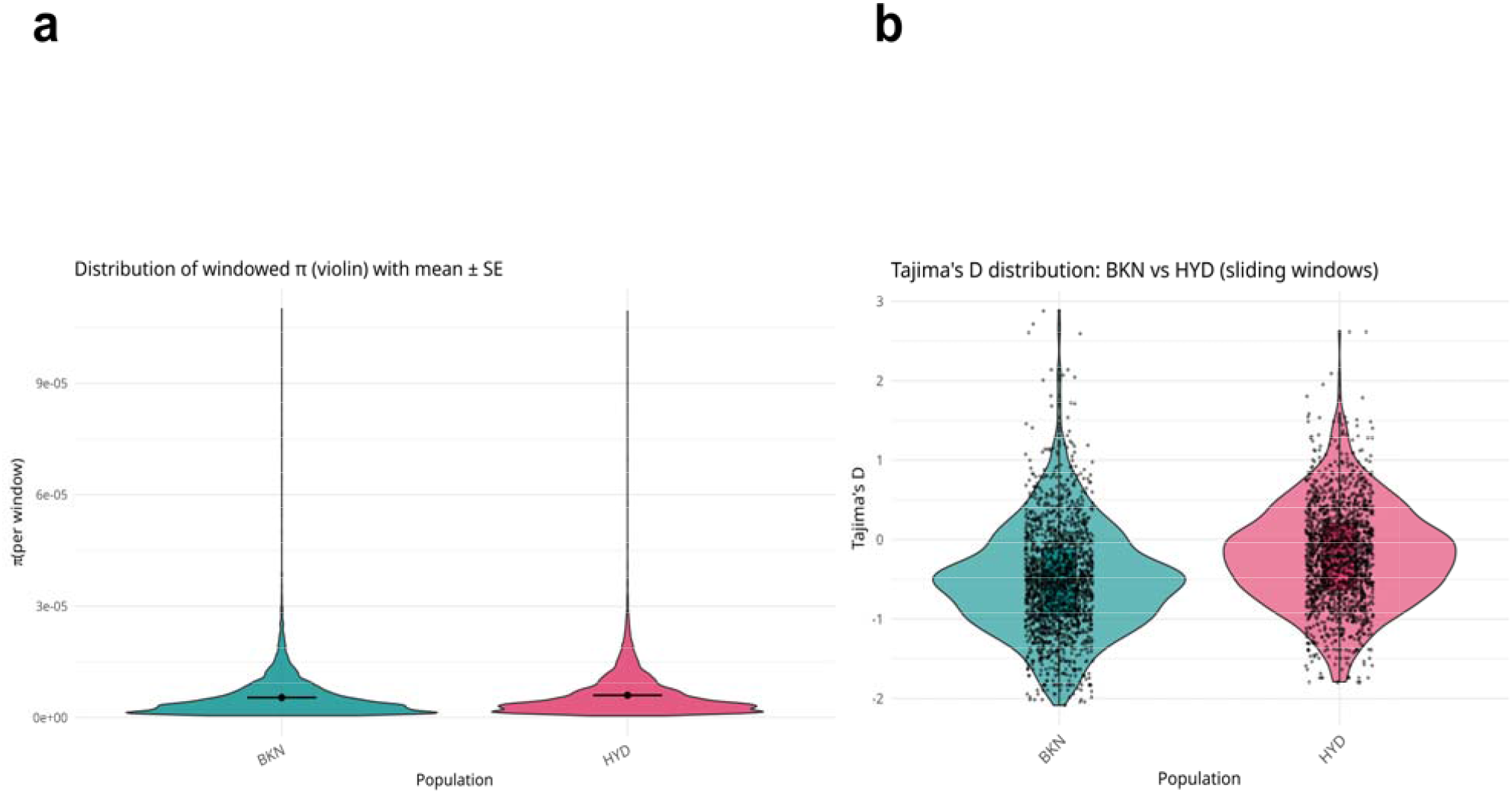
Genetic diversity estimates across population groups. (a) Windowed nucleotide diversity distribution. shown using violin plots for BKN (n=8,813 windows; mean π = 5.42×10^−6^, median = 3.70×10^−6^) and HYD (n = 8,408 windows; mean π = 6.05×10^−6^, median = 3.88×10^−6^). Both populations show very low genome-wide diversity consistent with a small long-term N_e_. **(b)** Violin-plots of **windowed Tajima’s D** (windows of 100 SNPs; see Methods) for BKN and HYD. Mean D value is negative in both populations (BKN mean = −0.51, HYD mean = −0.20), indicating reduced genetic diversity and a skewed SFS.

Although our sample size limits the Tajima’s D estimates, and no sweeping statements can be made about the frequency of the rare variants, nucleotide diversity estimates remain robust (mean π of the order of 10^−6^) and underscore low standing diversity due to small long-term effective population sizes (Ne) in both sites. *Refer to supplementary results section for a discussion of the diversity estimates in context of inbreeding results*.

### Relatively Higher Mutation Load in BKN is Driven by More Recent Inbreeding and Long-standing Small N_e_

The predictions from the theory of genetic drift for populations of small historical N_e_ imply that such populations accumulate deleterious mutations over time, which further present negative fitness consequences directly tied with extinction risk (Ohta 1973). Conversely, purifying selection eliminates these lethal variants and offers a fleeting reprieve. To understand the mutation-selection dynamics, we quantified per-sample load in our cohorts from 58,931 polarised, biallelic SNPs using a polarisation pipeline (see Methods). Individual level summaries highlight tight clustering of total derived-allele counts: : total derived alleles per sample were ≈74,500-76,500 900 alleles (≈1-1.5% relative width; see ***Supplementary table S9***). Individuals were also comparable across recessive load and SnpEff-based impact-stratified counts. Across samples, the per-sample median counts by impact were approximately: **HIGH** ≈ **9–10 homozygous-derived sites, MODERATE** ≈ **266–269, LOW** ≈ **680–690**, and **MODIFIER** ≈ **35**,**600** (per-sample count medians).

Inferring group statistics, the BKN individual had higher mutation load measures than HYD individuals (see ***Supplementary table, S10***). For BKN and HYD groups, the mean additive (burden) per individual were **2**,**618.5** (SD = 410.1) and **2**,**076.4** (SD = 363.9), the mean homozygous-derived counts (recessive load) were 37,365.7 (SD = 74.0) and 36,538.4 (SD = 39.0), and the mean heterozygous counts were 38,838.8 (SD = 71.5) and 38,069.6 (SD = 49.2), respectively.

These differences are visually reinforced in the group-level distributions, where BKN distributions are positioned upwards for both additive burden and homozygous-derived counts (***Fig. 6a***, *also see* ***Supplementary Fig. S15a-b***). Apart from the total mutation load at the individual level (***Fig. 6b***), the sample-level scatter shows a tight linear relationship between total derived alleles and homozygous-derived counts across samples (***Fig. 6c***), suggesting that samples with overall higher numbers of derived alleles also tend to harbour more homozygous-derived sites.

**Fig 6.**
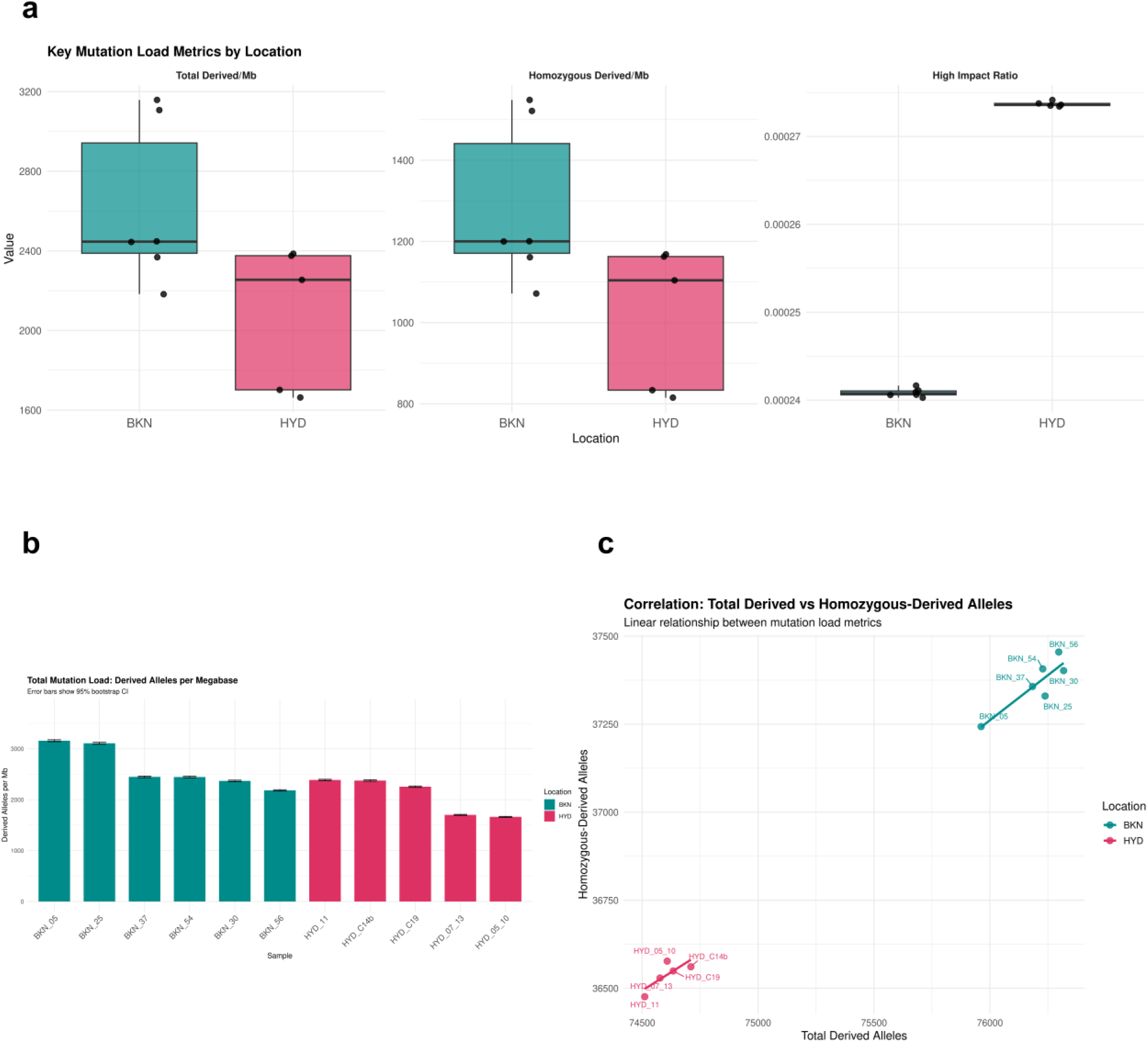
Individual and per-sample mutation-load metrics. **(a)** Boxplot of per-sample **derived-allele burden** (left), **homozygous-derived counts** (middle), and **high-impact ratio** (right) normalized by callable bases. **(b)** Bar-plot of per-sample **derived-allele counts per megabase** with 95% bootstrap CIs (samples ordered by location). **(c)** Scatter of **total derived alleles versus homozygous-derived** alleles showing a tight linear relationship and population separation.

The elevated counts of homozygous-derived alleles and additive burden are congruent with demographic processes that increase homozygosity - such as small Ne, recent bottlenecks, or elevated inbreeding - which improve the probability of derived alleles being exposed in the homozygous state (Charlesworth and Willis 2009; Hedrick and Garcia-Dorado 2016). However, the abundance of non-coding and low-impact consequences in the consequence profile indicates that much of the additional load in BKN is driven by low-effect or non-coding variants rather than an excess of strongly deleterious protein-truncating alleles. Additionally, we note that the overall mutation burden in HYD was positioned downwards mainly due to the higher callable sites in the merged samples (HYD_05_10, HYD_07_13). The functional consequence breakdown of the polarised polymorphisms confirms that most segregating sites align with noncoding categories, and protein-altering polymorphisms constitute a small minority (***Supplementary Fig. S15c***).

### Demographic Reconstruction Reveals Early Divergence and Prolonged Population Isolation

We reconstructed the recent demographic history of the two focal populations (BKN, HYD) from a folded two-population joint SFS and fitted four alternative models in ∂a∂i (SI, IM, AM, SC)(see *Methods*). Model comparison by AIC strongly favoured the **Ancient Migration (AM)** topology as the best description of the data (AIC values: AM = 481,226.4, IM = 523,576.3, SI = 529,306.7 and SC = 526,110.3 (ΔAIC≫10 in all pairwise comparisons), indicating the joint SFS is most compatible with an early divergence followed by a transient interval of post-split gene flow that ceased long before the present (see ***Supplementary Table S11*** for summary statistics for all model runs; refer to ***Supplementary Tables, S12 and S13*** for detailed breakdown for all models).

Parametric bootstrap replicates resulted in tightly concentrated bootstrap distributions and small standard deviations for the parameters in our runs, suggesting stable convergence under the fitted configuration and the information content of the folded joint-SFS for this dataset. Diagnostic plots (see ***Fig. 7a***) show that the AM model envisages the overall shape of the SFS with the largest residual values confined to the extreme corners of the matrix (low-frequency cells), suggesting a generally good fit across frequency classes.

**Fig 7.**
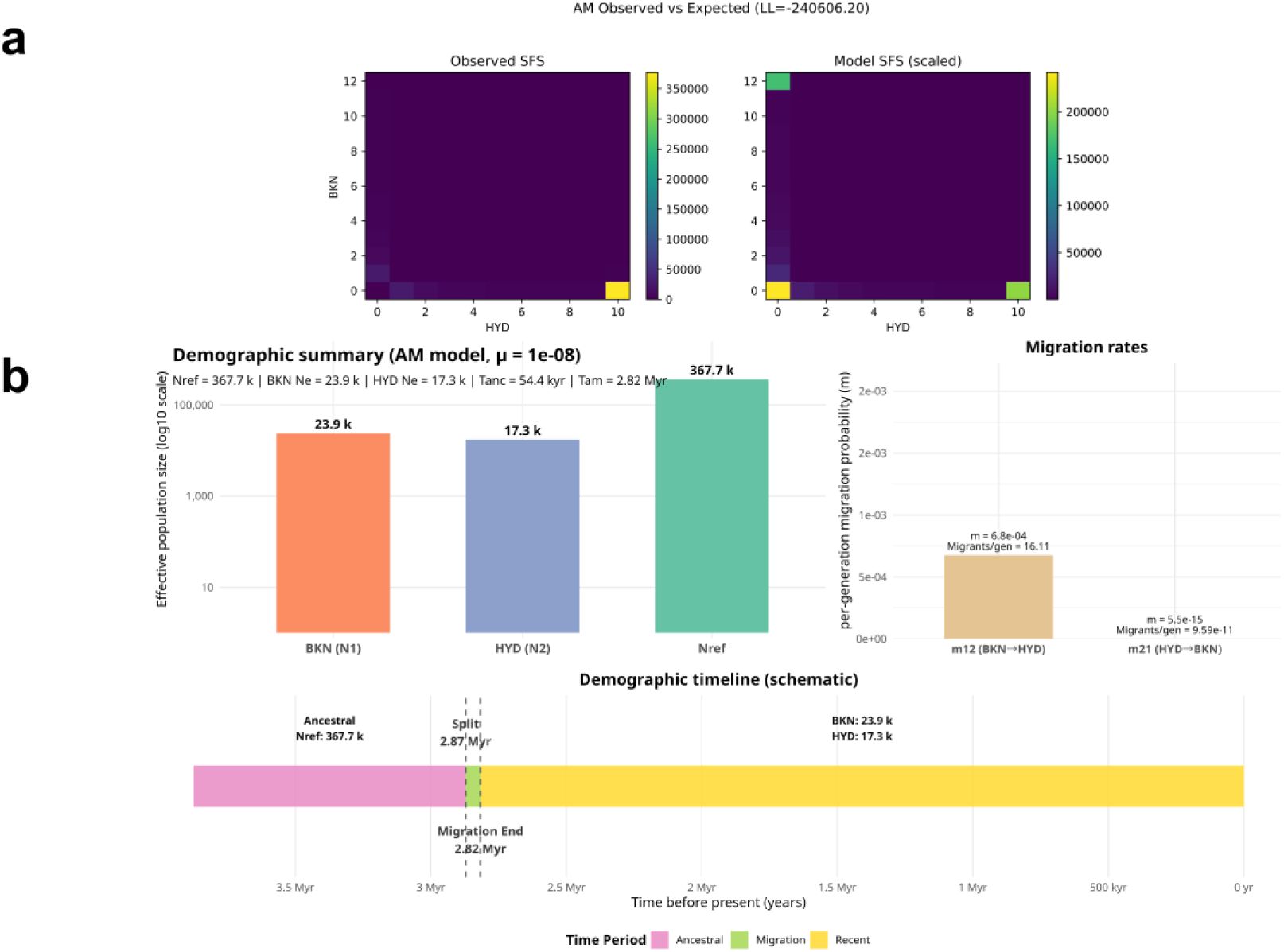
SFS fitting and Demographic timeline inferred under the AM model. **(a)** Top left: folded two-population joint site-frequency spectrum (BKN × HYD) used for demographic inference. Top right: corresponding model-predicted SFS under the best-supported Ancient Migration (AM) topology (scaled to sample sizes). Panels **(b)** and **(c)** show the inferred demographic histories for BKN and HYD species population with parameter conversions to absolute time assuming g = 12 yr and alternative mutation-rate calibrations. Both reconstructions support an early split followed by a short interval of asymmetric gene flow into BKN and prolonged isolation thereafter, along with a precipitous post-split reduction in N_e_. Parameter uncertainty was assessed by parametric bootstrap; model comparison (AIC) strongly favours the AM topology over SI/IM/SC/ (ΔAIC>>10).

Interpreting demographic times and effective sizes requires a standard choice of mutation rate (***µ***) and generation time (g). When using a composite generation time (g) of 12 years and ***µ*** in the range common for avian genomes (1-2 × 10^−8^ per site per generation), the qualitative portrait is maintained while absolute timescales shift linearly (Robinson 2022). Under ***µ*** = 2 × 10^−8^, the inferred split and end-of-migration map to roughly ≈1.4–1.5 million years ago (split ≈1.44 Myr; migration ended ≈1.41 Myr; see ***Supplementary Fig. S16b***). With ***µ*** = 1 × 10^−8^, these dates roughly double (***Fig. 7b***), illustrating sensitivity to mutation rates but preserving the qualitative conclusion: an early divergence followed by a fleeting ancient window of gene flow and subsequent isolation.

The relative population sizes (nu1 and nu2) **indicate that both extant populations are sharply reduced relative to the inferred ancestral population**. Scaled N_e_ (using **θ** and reasonable ***µ*** values) **places contemporary N**_**e**_ **for BKN and HYD vulture species in the low (10**^**3**^**-10**^**4**^) range in our conversions (≈8 to 24 k; ***Fig. 7*** and ***Supplementary Fig. S16***). In contrast, the ancestral N_e_ is substantially greater, consistent with a genome-wide reduction in the effective population size post divergence. *Supplementary section further underscores the methodological caveats associated with the parameter estimation*.

## Discussion

The combined genomic picture that emerges from our analyses (ROH, heterozygosity, π, population structure, and polarised mutation-load counts) coherently illuminates a two-stage demographic process with predictable genomic consequences. We find that vulture species sampled in both locations inhabit the hallmarks of long-standing reductions in effective population size (Ne) and recent demographic isolation, although BKN samples show somewhat stronger recent drift and inbreeding, accompanied by a conspicuous rise in realised mutation burden. These trends contrast with the actual population decline in the Indian subcontinent, which started in the 1990s with a dramatic increase in Diclofenac use in animal husbandry (SoIB 2023).

The F_*ROH*_ values reported place the most inbred vultures in the same order of magnitude as some of the extreme, well-documented bottlenecked examples from the literature. For instance, late Wrangel-Island woolly mammoths have been described with very low heterozygosity and long ROH presenting F_*ROH*_ often in the 0.3-0.5 range (Dehasque 2024; Palkopoulou 2015). Isle Royale wolf populations likewise exhibit very high ROH and F_*ROH*_ in the 0.2-0.5 span in the most bottlenecked islands (Robinson 2019). The estimates also approach those reported for other threatened/endangered mammals and birds -- for example Indian tigers (Khan 2021), Florida panthers (Saremi 2019) and Montane red foxes in North America (Quinn et al. 2024) -- but remain substantially lower than the extreme autozygosity documented for the Critically Endangered Chatham Island black robin which displayed F_*ROH*_ as high as 0.63 in some populations (Seth 2022).

Instead of interpreting each metric in isolation, the data combine to communicate a temporally layered process, with tales of demographic history and inbreeding spanning most of the entirety. Historical population size (Ne) reductions— reflected in low nucleotide diversity (π) — likely dampened the efficacy of purifying selection and lifted the baseline of weakly deleterious variation segregating in the populations (Ceballos et al. 2018a; McQuillan 2008; Xue 2015). Superimposed on that background, recent demographic constriction or consanguinity, as evidenced by the presence of long ROH tracts and elevated F_*ROH*_ values, shows that these homozygous tracts disproportionately reveal derived alleles and thereby inflate realised recessive load (Charlesworth and Willis 2009; Hedrick and Garcia-Dorado 2016). The results of mutation-load quantification further reinforce this understanding: BKN, as compared to HYD, is represented by a relatively higher additive burden and more homozygous-derived sites primarily attributable to LOW-impact/noncoding categories. This pattern is congruent with population-genetic theory predictions when N_e_ plummets: the efficacy of purifying selection subsides for mutations of small effect size, so they drift to higher frequency and can become fixed as homozygous, while highly deleterious alleles are efficiently expunged or remain sporadic. Therefore, the observed burden in BKN is construed as an aggregate effect of many mildly deleterious variants being exposed by the actions of drift/inbreeding, rather than a proliferation of high-impact alleles (Henn et al. 2015; Keightley and Eyre-Walker 2007; Simons et al. 2014).

The functional dissection of the mutations still has meaningful biological implications. Because most additional homozygous derived sites in BKN are driven by low-impact or regulatory mutations, their discrete effects are small; yet when many such sites are homozygous throughout the genome, their polygenic load can depress fitness components (fecundity, longevity, immune competence) that arise from many loci acting jointly (Keightley and Eyre-Walker 2007; Simons and Sella 2016; Simons et al. 2014). A thorough empirical comparison across taxa reveals that such small, inbred populations frequently harbour elevated deleterious load that translates to lower fitness (e.g., Isle Royale wolves, Florida panthers, mountain gorillas), though the magnitude pivots on dominance coefficients and the extent of purging, if any (Robinson 2019; Saremi 2019; Xue 2015).

The site-frequency spectrum adds more nuance to this portrait. The low π reconciled with demographic interpretations indicates either a recent bottleneck followed by limited recovery or ongoing purifying selection on linked markers — a process that concentrates homozygosity via genetic drift (Li and Durbin 2011; Nielsen 2005). The demographic model that best explains the observed SFS (Ancient Migration with asymmetric early gene flow) further elucidates how subsequent isolation and drift contributed to the present-day divergence: a transient stream of gene flow succeeded by prolonged separation creates a genomic landscape with pockets of shared ancestry but an overall signal of reduced contemporary N_e_ (∂a∂i results, see Methods).

Another important clarification following from the structure analyses is that BKN grouping largely involving a single, internally coherent genetic cluster and an outlier represents two different taxa, supported by the field records at the site. PCA, ADMIXTURE, and the similarity-network results all show the BKN cluster tightly connected (very low within-site pairwise FST and high genetic similarity; ***Fig. 5; Fig. 6***), and the sole BKN outlier (BKN_UV_o_01) routinely projects away from that tight cluster, often toward HYD samples at higher resolution. Cumulatively, these patterns suggest one of the two simple explanations for the observed data: (i) the BKN panel is predominantly a single species (either Egyptian or Eurasian griffon), or (ii) the samples are a genuine mixture, but one species dominates the sampled genetic signal to the extent that it masks a minority lineage. Furthermore, our demographic inferences (Ancient migration with a large drop in the BKN lineage, ***Fig.7***) and region-wide monitoring show dramatic declines for Egyptian and griffon vultures when using a 2000s baseline, which would make genomic recovery at BKN challenging for any rare species (SoIB 2023). **In essence, the clearest mechanistic reading is that both sites have experienced sustained drift, with BKN vulture species more strongly affected by recent isolation or founder effects apparent in the higher realised burden of derived alleles**.

## Conservation Interpretation with Methodological Caveats

Although our dataset provides coherent genome-wide signals of low diversity, elevated autozygosity, and increased mutation burden, these patterns must be interpreted in light of the methodological constraints inherent to reduced-representation sequencing. The pilot-run curation steps were essential to minimize artefactual structure and genotype distortion: BKN_24 was removed due to extreme missingness, BKN_35 was reclassified as an outlier individual, and two biologically duplicate pairs were merged at the read level to avoid redundant sampling while improving effective coverage. These steps resulted in improved downstream genotype calling and are discussed in detail in the Supplementary Results (***Supplementary Fig. S8–S10***).

Even after curation, ddRADseq surveys only a fraction of the genome, so summary statistics such as Tajima’s D, π, and heterozygosity-based F estimates remain sensitive to ascertainment bias, missing data, and locus representation (Nielsen 2005). Accordingly, we interpret the negative Tajima’s D and very low π as consistent with recent demographic contraction and/or purifying selection, rather than as exact estimates of demographic magnitude (Nielsen 2005; Tajima 1989). Likewise, the discrepancy between PLINK-based inbreeding coefficients and ROH-derived FROH reflects a known difference in sensitivity: the former is strongly affected by SNP composition and missingness, whereas the latter provides a more direct readout of autozygosity and recent inbreeding (Ceballos et al. 2018a,b; McQuillan 2008; Purcell 2007).

From a conservation perspective, the combined evidence for high ROH burden, reduced nucleotide diversity, and elevated mutation load is worrying because it indicates limited adaptive buffering and a potentially increased risk of inbreeding depression (Kardos et al. 2016, Hedrick and Garcia-Dorado 2016; Keller and Waller 2002). However, the exact severity of these risks should be viewed as conservative approximations, particularly because site-level comparisons are also influenced by species composition, founder history, and sample size. We therefore encourage readers to consult the Supplementary Results for the full curation workflow and supporting analyses, which provide the necessary context for interpreting the main-text conclusions.

## Implications for Conservation

Our genomic results — high F_*ROH*_, depleted π, and a skewed SFS and a higher realised mutation burden in BKN — imply more than an academic signal of the past decline. They directly translate into elevated extinction risk and invite a robust conservation framework addressing extrinsic threats and intrinsic genomic fragility we document.

Through the lens of conservation genomics, the synthesis of the demographic and functional signals points to an integrated conservation roadmap-

### a. Reduce extrinsic mortality

Maintaining and strengthening non-genetic protections (e.g., carcass-management practices) will reduce further N_e_ crashes and buy time for genetic intervention (Oaks 2004). Policy enforcement to eliminate veterinary diclofenac, promotion of vulture-safe alternatives (e.g., meloxicam), and continued monitoring of carcass residues are therefore non-negotiable conservation actions.

### b. Institute genomics-informed captive management

Higher realised recessive mutation burden in the more drift-impacted cohort suggests two related management priorities: (a) reduce immediate mortality so small populations can grow (lowering the per-generation strength of drift), and (b) implement genomics-informed breeding and release schemes where captive breeding is used (e.g., breeding centres in India; Wildlife Institute of India 2016). The other practical steps include expanding spatial sampling to capture within- and among-site species composition and combining ddRAD panels with targeted whole-genome resequencing for founders and potential donor populations used for augmentation and other critical decisions.

### c. Controlled genetic augmentation

Although genetic rescue when outbreeding risk is low yields large, rapid fitness gains, for vultures, augmentation (carefully sourced translocations) can only be considered after thorough risk assessment (disease, outbreeding depression, species composition, local adaptation, genetic matching to minimise maladaptation and pathogen induction) and with robust genomic monitoring to evaluate impacts. (Frankham 2015; Frankham 2016; Hedrick and Garcia-Dorado 2016). It is equally imperative to embed genomic sampling into routine demographic and health monitoring so that changes in F_*ROH*_, π or mutation-load can be tracked alongside survival, reproduction, and disease outcomes.

### d. Link genomics to phenotype

Prioritising targeted functional assays (transcriptomics, immunocompetence, toxicology, reproductive success) can help establish whether the genomic load manifests as measurable fitness deficits (survival, reproduction, disease resistance) — these empirical links are indispensable to assess the benefits and risks of actions such as genetic rescue (Henn et al. 2015; Simons et al. 2014).

Finally, social and operational realities must also be addressed. Sustained funding, supply chains for safe drugs, community engagement (carcass management and vulture safe zones), and coordination with other agencies (environment, animal husbandry, public health) are required to translate genomic cues into demographic recovery. The breeding centres also face lesser documented challenges: small founder sizes, breeding failures, behavioral variations in captive-bred birds, and the logistical complexity of establishing Vulture Release Areas must also be addressed explicitly in national and state action plans. At its core, genomic actionability is high — but it must be embedded in sustained threat reduction, veterinary policy enforcement, and rigorous health and ecological risk assessment to achieve tangible conservation outcomes.

## Resource Availability

### Data Availability

Raw dd-RAD sequencing data of the 15 newly generated vulture genomes have been deposited at the NCBI-SRA database with BioProject accession PRJNA1404108 and will be made publicly available after the set release date.

### Code and Script Availability

The analysis code and scripts used for ddRAD data processing and population-genetic analyses are available at: Shukla, M. (2026). *Indian Vulture Population Genomics (ddRAD-data processing and analysis)* (Version v1.0) [code]. Zenodo. DOI: https://doi.org/10.5281/zenodo.18331891.

> Project name: Indian Vulture Population Genomics (ddRADseq data processing and analysis)
>
> Project home page: https://github.com/ShuklaManas578/Indian-Vulture-Population-Genomics-ddRAD-data-processing-and-analysis-
>
> Operating system(s): Linux
>
> Programming language: Python (3.10.19) and R (4.5.0)
>
> License: NA

### Funding

This work received no financial support from any funding agency for the research and publication of this article. The ddRAD-seq genotyping was partially supported by an intramural grant from the Institute of Eminence (IoE), University of Hyderabad, and Department of Biotechnology (Govt. of India) funded BUILDER grant.

### Authors’ Contributions

MS (Conceptualization [Lead], Analysis, Methodology [lead], Visualization, Writing—original draft [lead], Writing—review & editing [lead], Software [Lead]), DLB (Writing—review & editing [supporting], Resources), BR (Resources), LN (Resources), SK (Methodology [Supporting], Writing—review & editing [Supporting], Supervision), VT (Conceptualization [Lead], Methodology [Supporting], Writing—original draft [Supporting], Writing—review & editing [Lead], Software [Supporting], Supervision).

## Acknowledgements

The authors acknowledge the Department of Systems and Computational Biology at the University of Hyderabad for lending the lab facilities for the wet-lab experiments. The authors would further like to extend their gratitude to the state forest departments and all the staff at the VCBCs for their work in the collection of the samples, especially Mr Shakeel. We also thank the director and curator at Nehru Zoological Park, Dr Sunil S. Hiremath and Ms J. Vasantha, respectively, for their help in obtaining the sampling permissions.

## Supplemental Materials

### Supplementary Figure Captions

**Fig S1.**
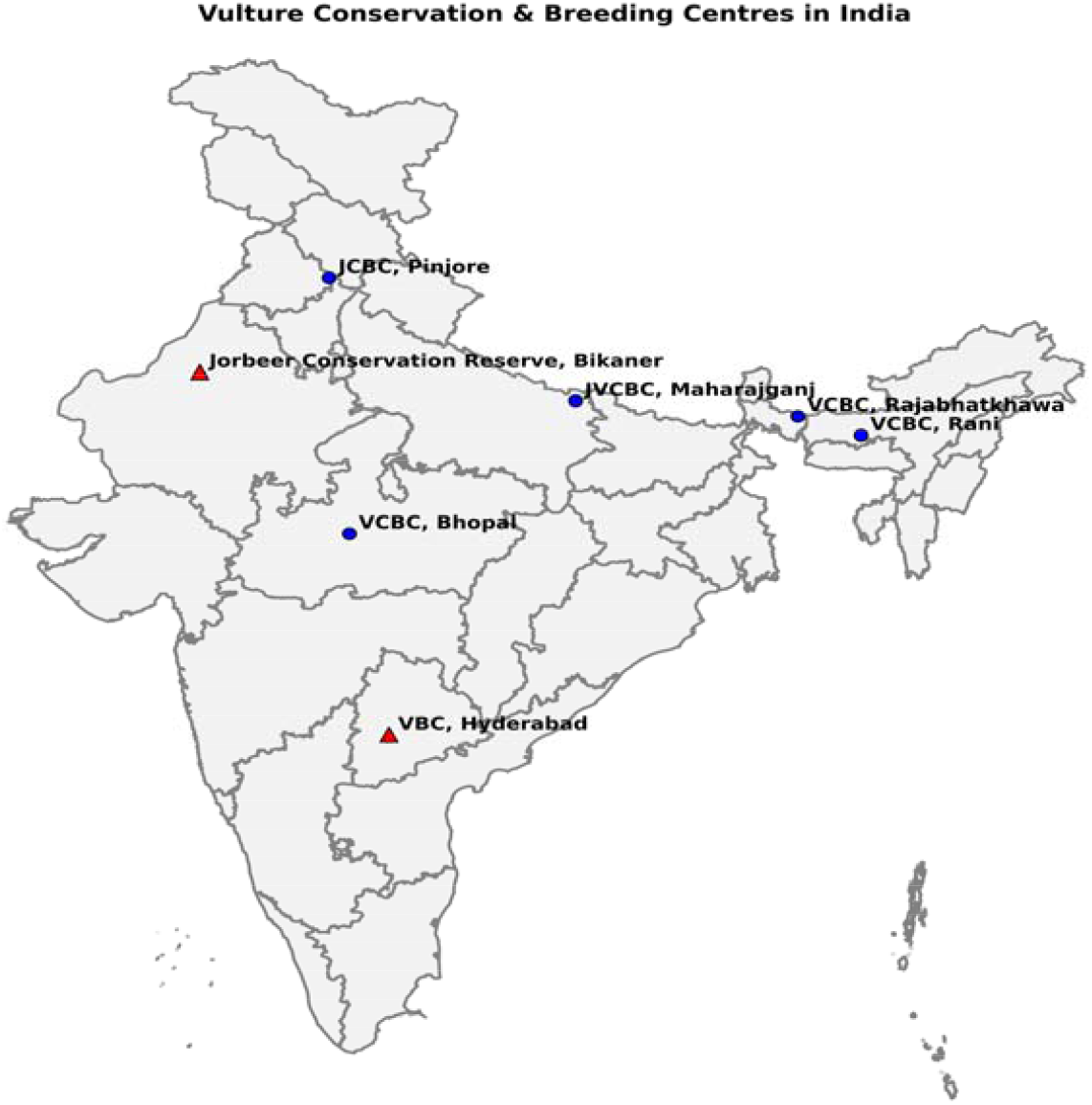
Vulture breeding and/or conservation centres in India. Red triangles represent the sampling sites while the blue circles indicate other breeding centres. The map was generated using a custom python script that used a shapefile (.shp) borrowed from the github page.

**Fig S2.**
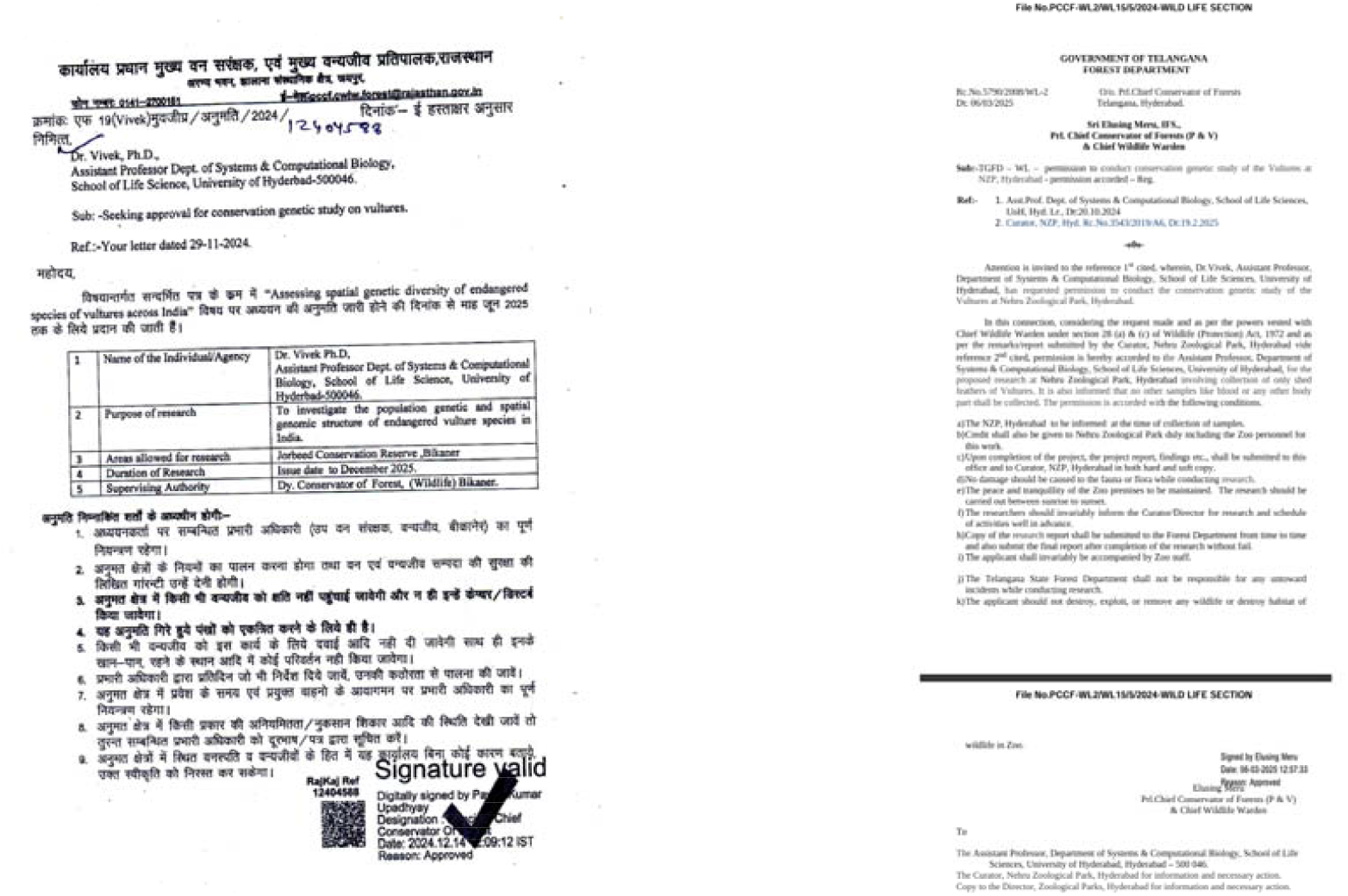
Sampling Permissions for the study. Permissions from state departments executives **(a)** PCCF, Rajasthan and **(b)** PCCF, Telangana enabling the collection of moulted feathers as biological samples for the duration of this study.

**Fig S3:**
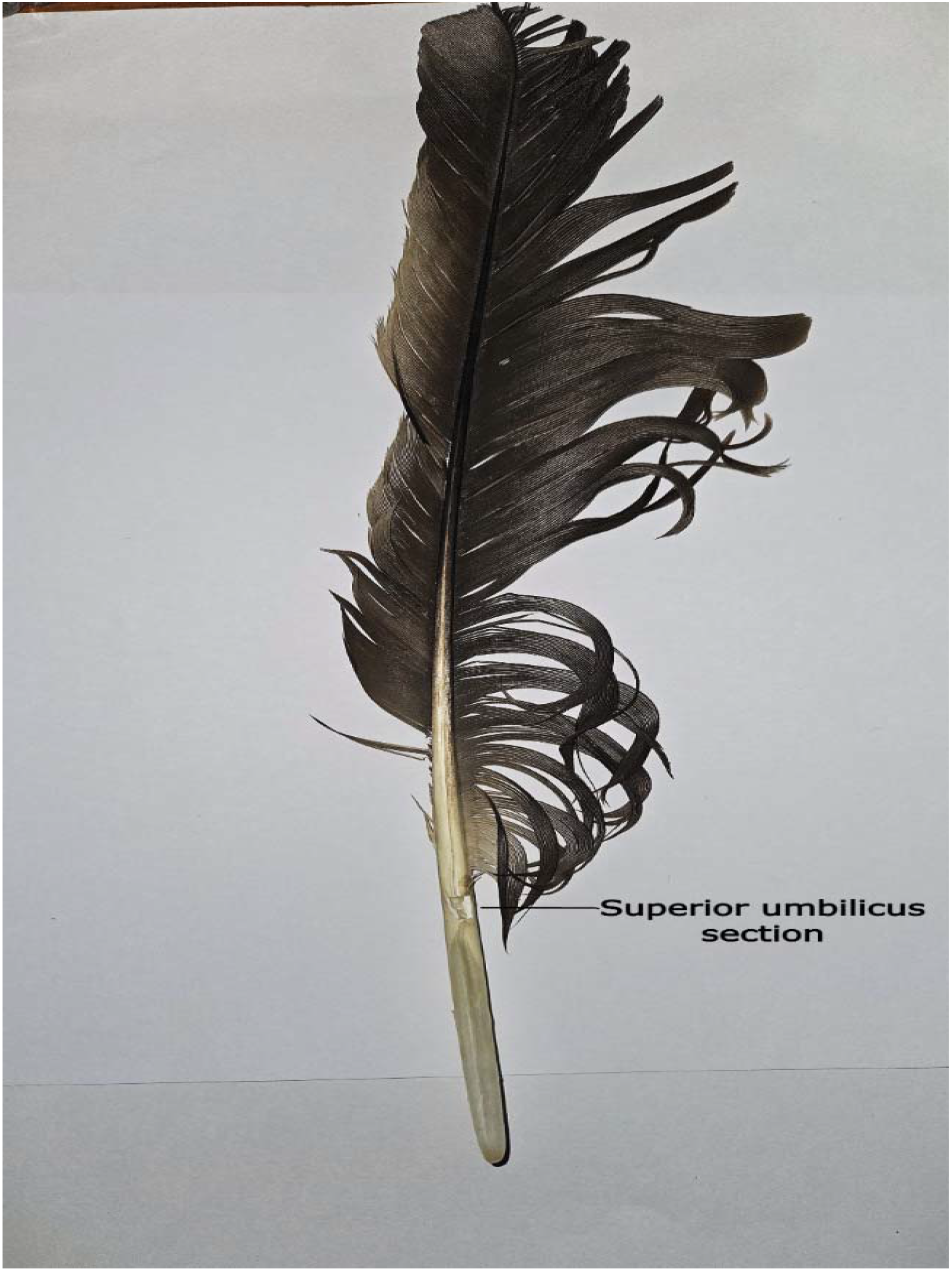
Retrix showing the superior umbilicus region. A retrix (tail feather) belonging to *Gyps bengalensis* showing the region of superior umbilicus post sectioning. A dried clot over the region was used to extract DNA from all the feathers.

**Fig S4:**
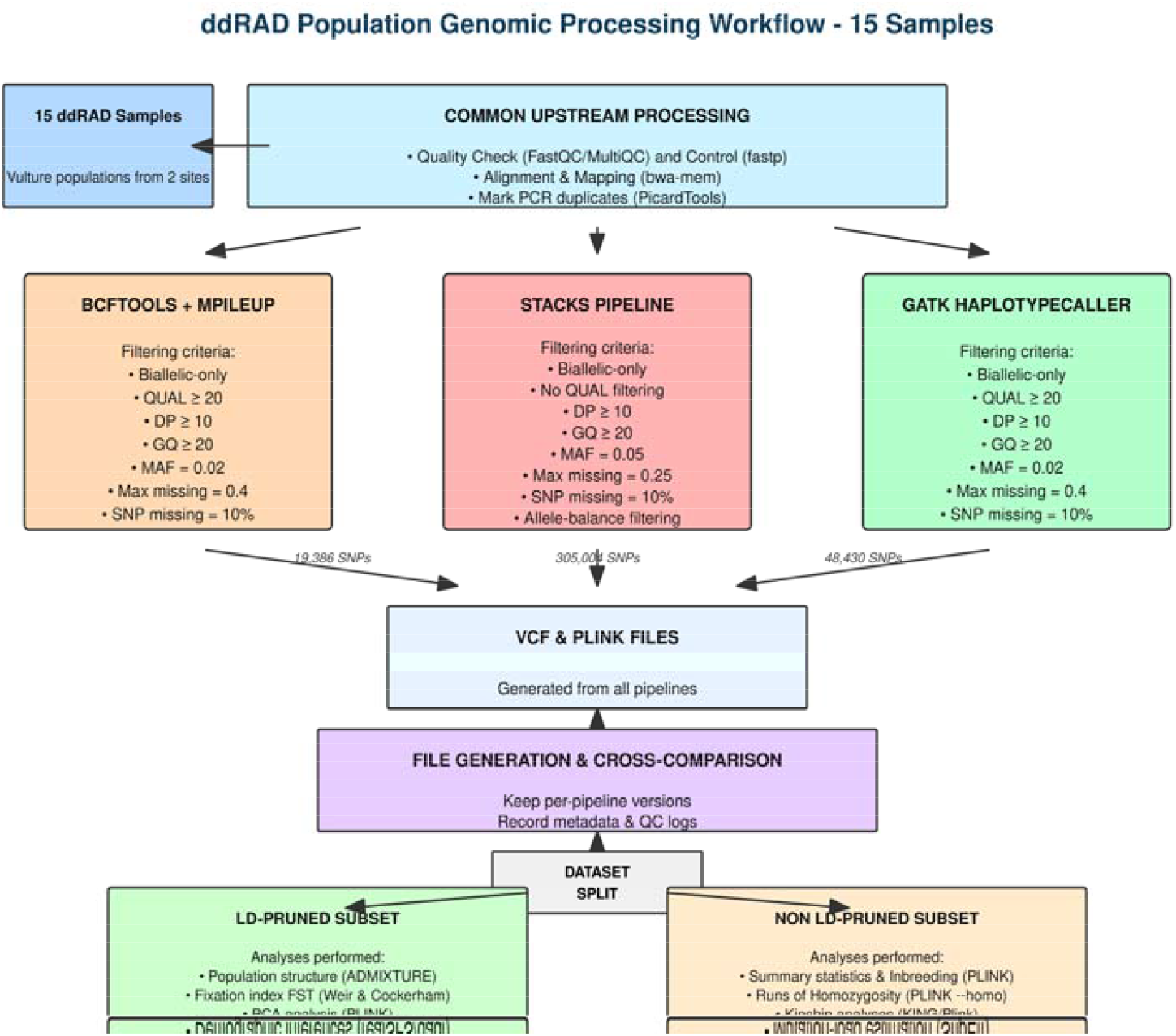
Pilot ddRAD-seq Data Analysis Workflow. The figure shows the preliminary analysis pipeline used to call SNPs and check for data quality issues flagging possible duplicates, and identifying group outliers.

**Fig S5.**
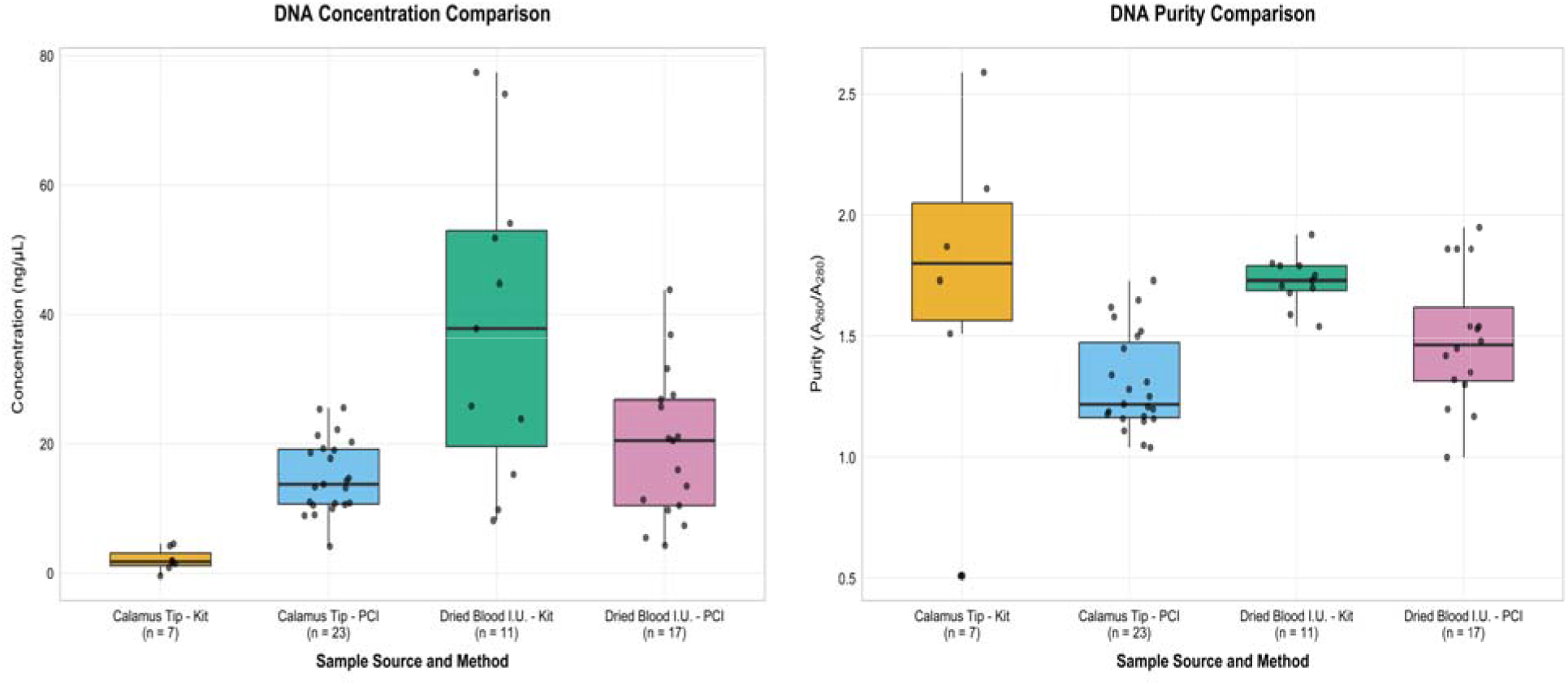
DNA yield and extraction consistency across protocol and sample types. The left panel shows DNA concentration (ng/μL) across sample and protocol types while the right panel exhibits DNA purity in terms of protein and phenol contamination. (I.U. stands for Inferior umbilical clot.

**Fig S6.**
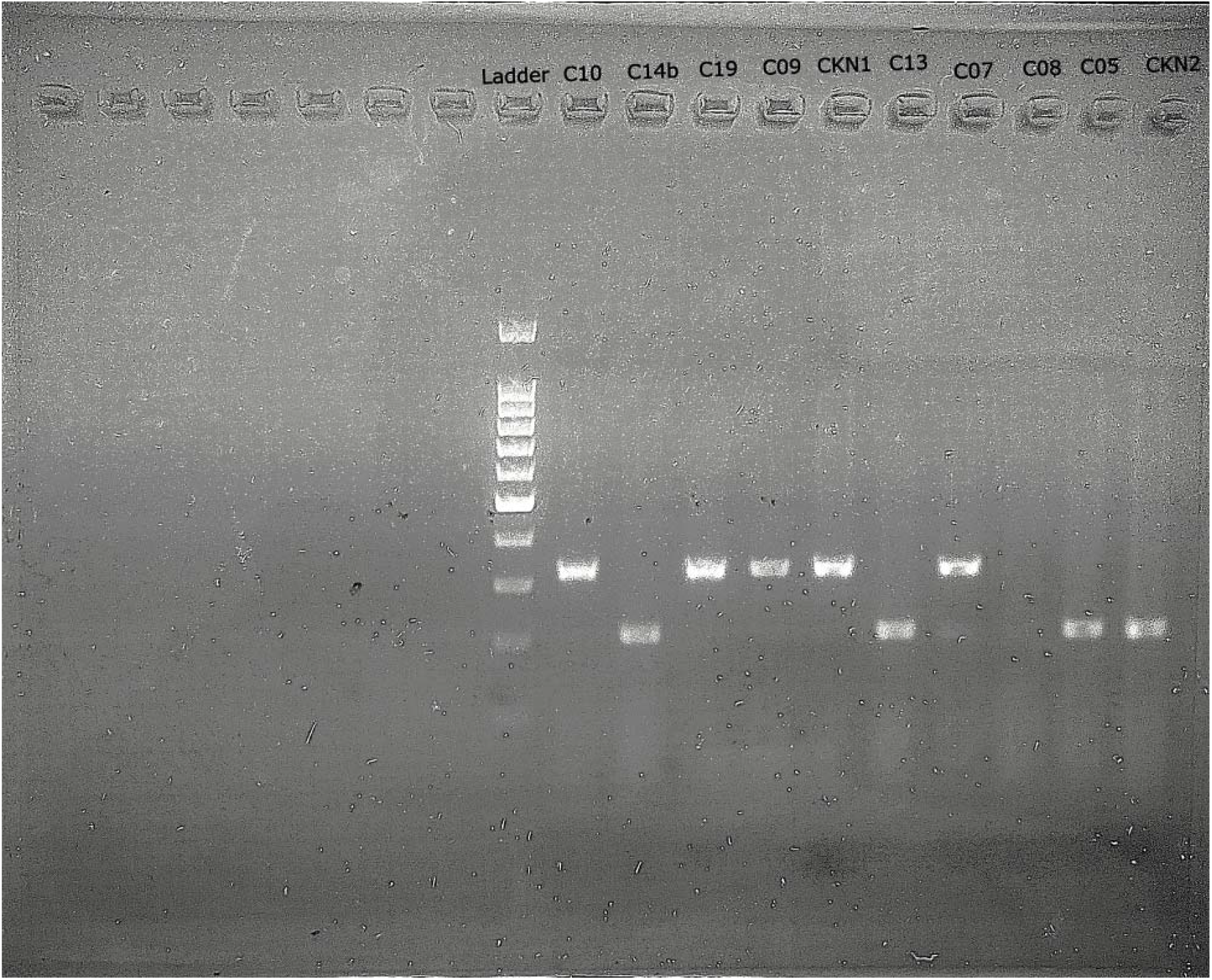
Gel electrophoresis of the DNA amplified from the vulture samples. The figure shows two bright bands corresponding to the product sizes of 322 and 210 bps for primer pairs 1 and 2, respectively. Vulture samples have a prefix ‘C’; while chicken samples have a prefix ‘CKN’, which was used as positive control for both the primer set reactions. The sample C08 did not show a band and was dropped in the primary QC analysis.

**Fig S7.**
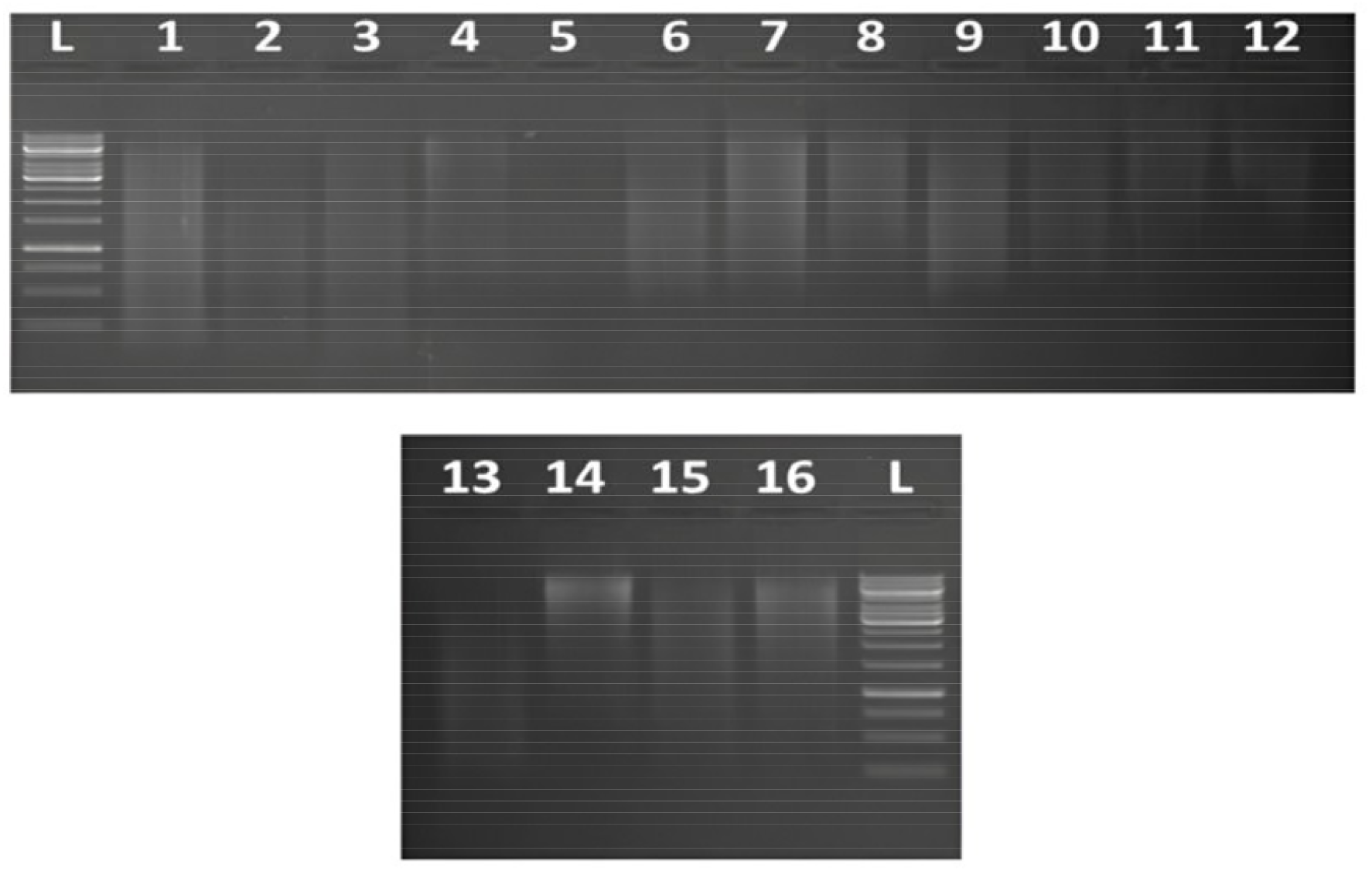
DNA QC by agarose gel electrophoresis. 1% gel electrophoresis results captured on Vilber Bio-Print Documentation system, where the lanes from 1-16 correspond to samples shown in the same order as in *supplementary table S3*.

**Fig S8.**
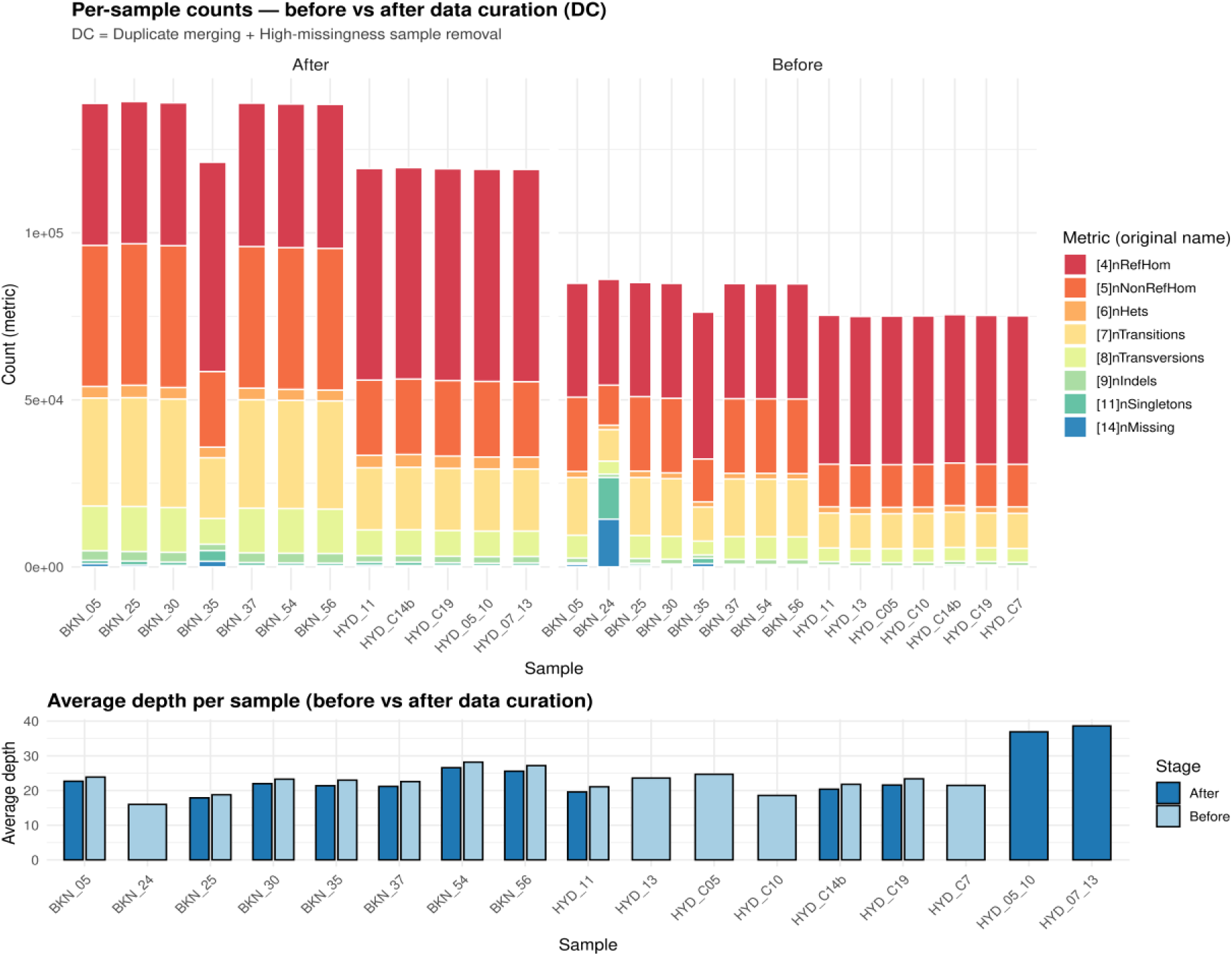
Genotype breakdown and depth comparison before and after merging duplicate samples. Figure shows the genotype calls before and after data curation which involved merging duplicate samples (*WR_05 + WR_10 and WR_07 + WR_13*) while simultaneously excluding the sample with high missing genotype calls (BKN_24). The top panel shows breakdown of the SNPs called for a number of metrics while the bottom panel compares average depth per sample before and after performing data curation.

**Fig 9.**
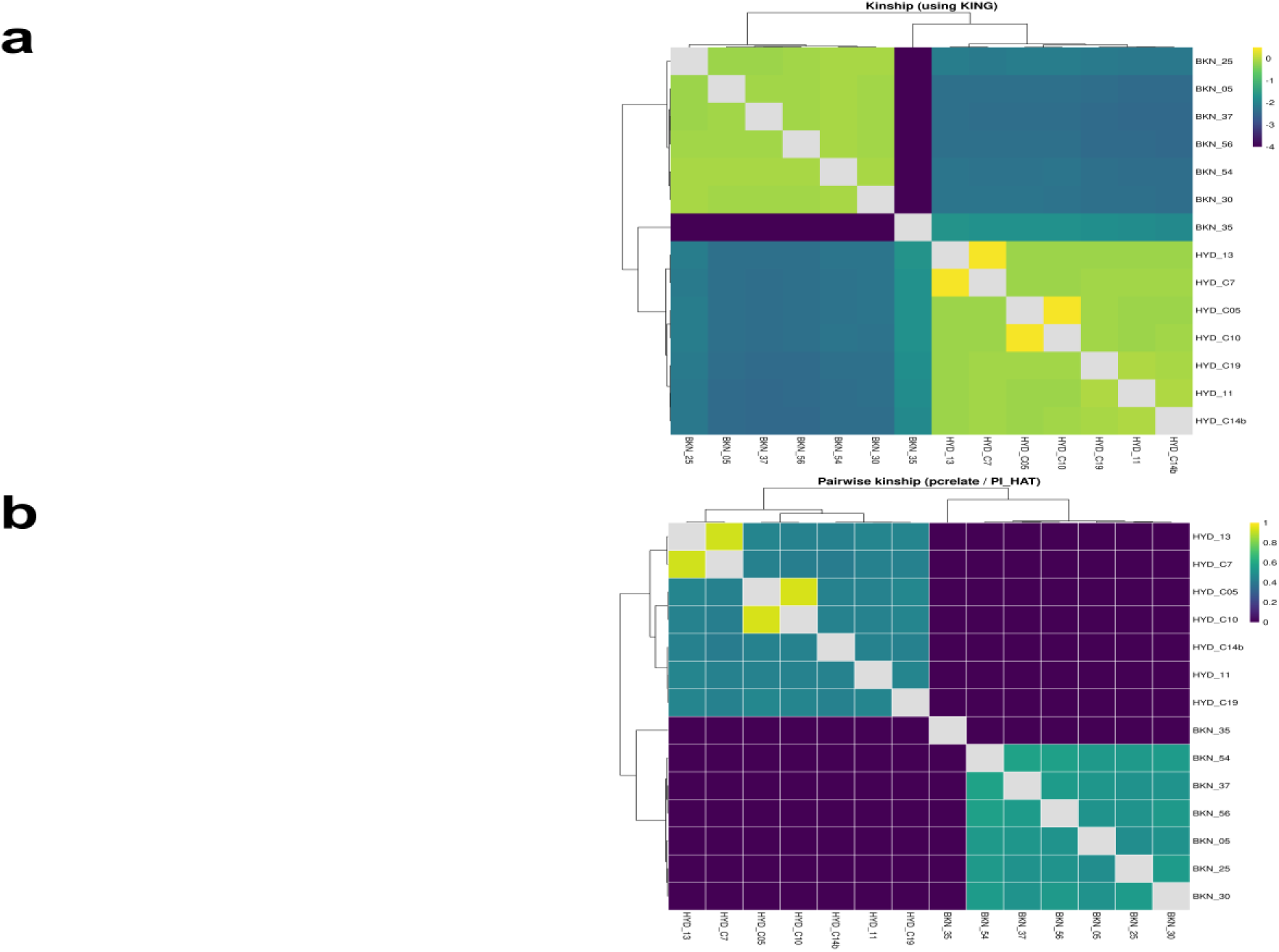
Kinship analysis and sample relatedness. **(a)** KING (tuned to vultures) identified two possible duplicate pairs shown in the yellow correlation squares: i)-WR_05 + WR_10, and ii)-WR_07 + WR_13. The duplicate samples were merged to improve effective coverage for those samples as already documented in past empirical works. The findings were in agreement with the results obtained using PLINK’s piHAT computation (*see supplementary* ***Fig. S6***). KING thresholds: >= 0.354 : Duplicate/MZ; 0.177 - 0.353 : 1st-degree; 0.0884 - 0.1769 : 2nd-degree; 0.0442 - 0.0883 : 3rd-degree; < 0.0442 : Unrelated (Manichaikul et al. 2010). **(b)** The kinship matrix generated using PC-Relate recapitulated the findings from KING’s relatedness analysis: Samples i)-WR_05 & WR_10, and ii)-WR_07 & WR_13 were confirmed to be the two duplicate pairs (higher pairwise values correspond to higher overall relatedness between the samples).

**Fig S10.**
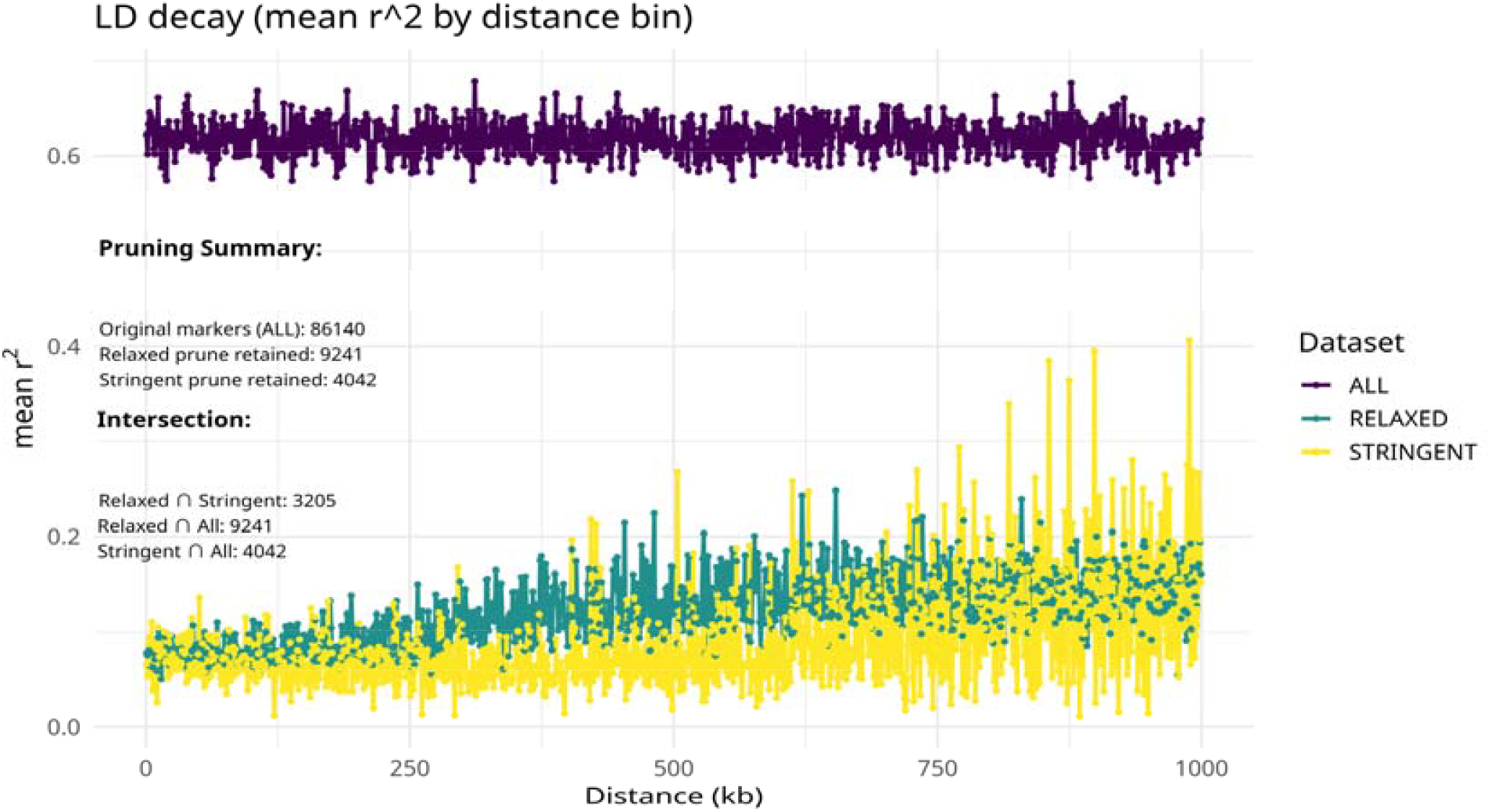
LD decay (mean r^2^) by physical distance. Mean linkage disequilibrium plotted against inter-marker distance (kb) for the full SNP set (ALL) and LD-pruned sets (RELAXED, STRINGENT). The pruning summary is included in the figure.

**Fig S11:**
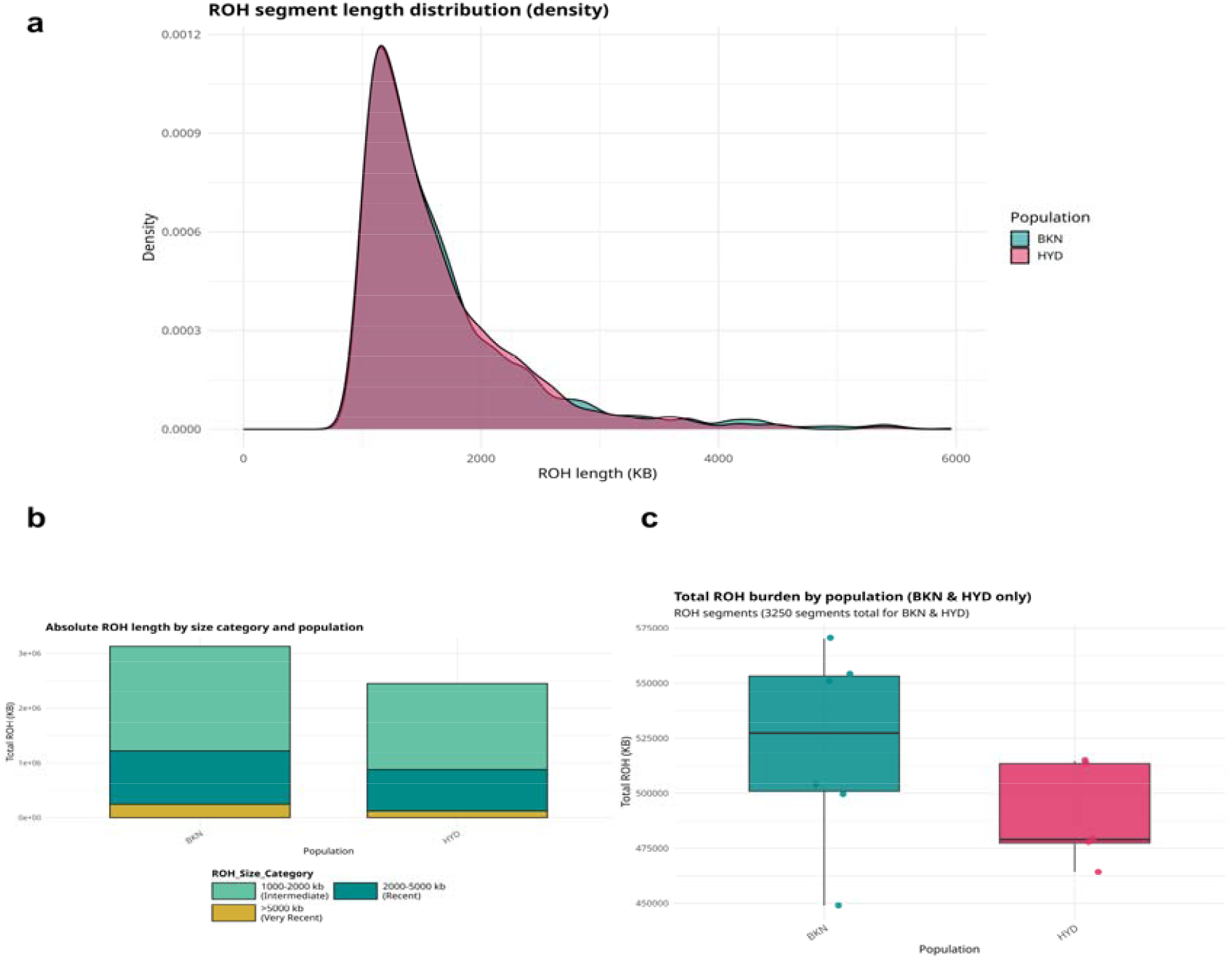
ROH and genomic autozygosity by population. **(a) Kernel density of ROH segment length**s (kb) pooled across individuals in BKN (teal) and HYD (rose); most ROH fall in the ∼1-2 Mb range with long tails corresponding to greater length. **(b) Proportion of total ROH** by size category for each population, depicting the relative contribution of recent (long) versus older (short) ROH classes. **(c) Group-level F**_***ROH***_ boxplots showing higher median autozygosity in BKN than HYD, consistent with elevated recent inbreeding in the BKN cohort.

**Fig S12:**
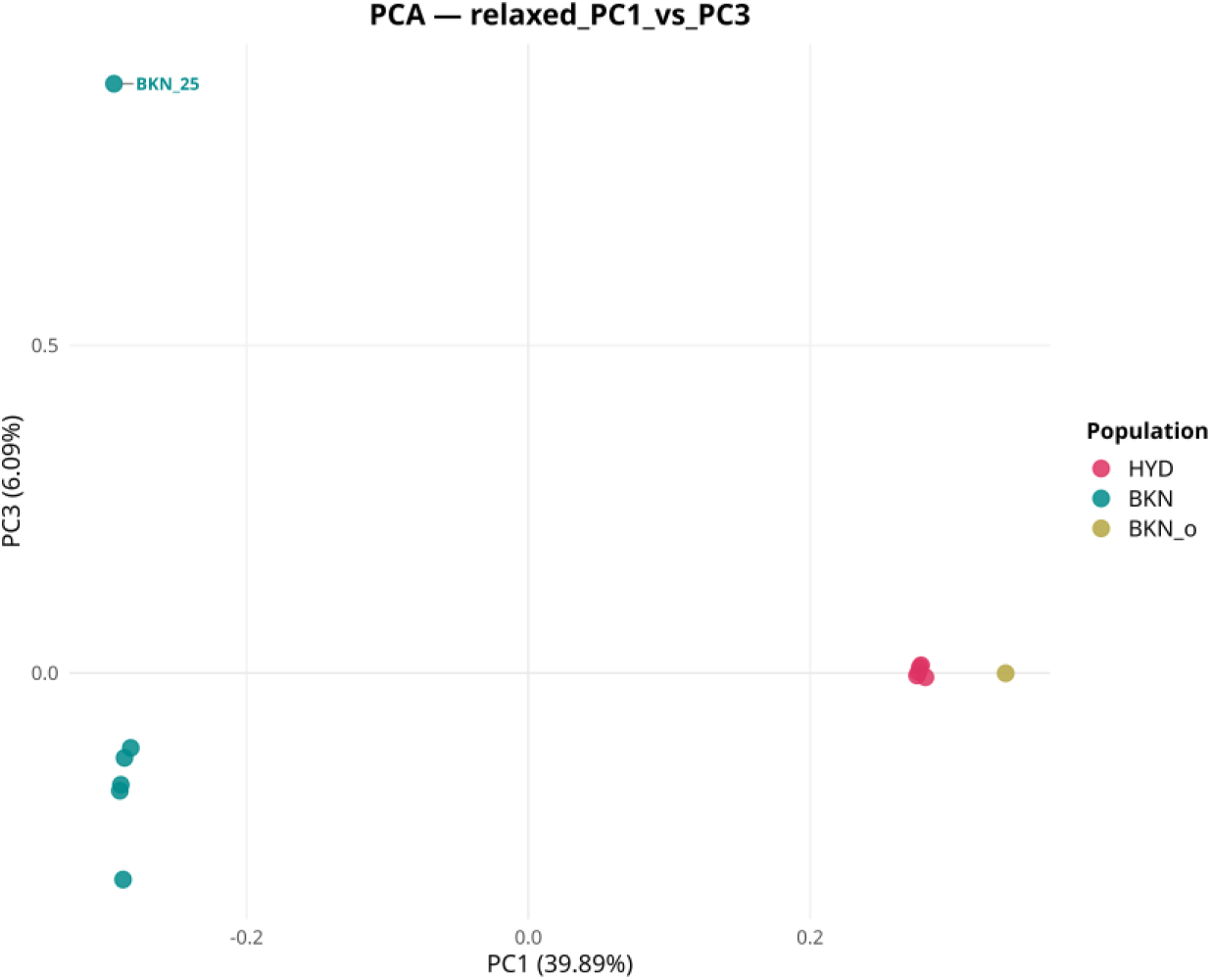
Principal Component Analysis. PCA of 12 samples (LD-pruned ‘relaxed’ SNP-set used) comparing the clusters in PC1 (39.9% variance) versus PC3 (6.09% variance) space. The grouping is in agreement with the results of ADMIXTURE and genetic similarity networks at higher resolutions, which also indicate partial affinity or distinct ancestry of the outlier sample to the HYD cluster.

**Fig S13:**
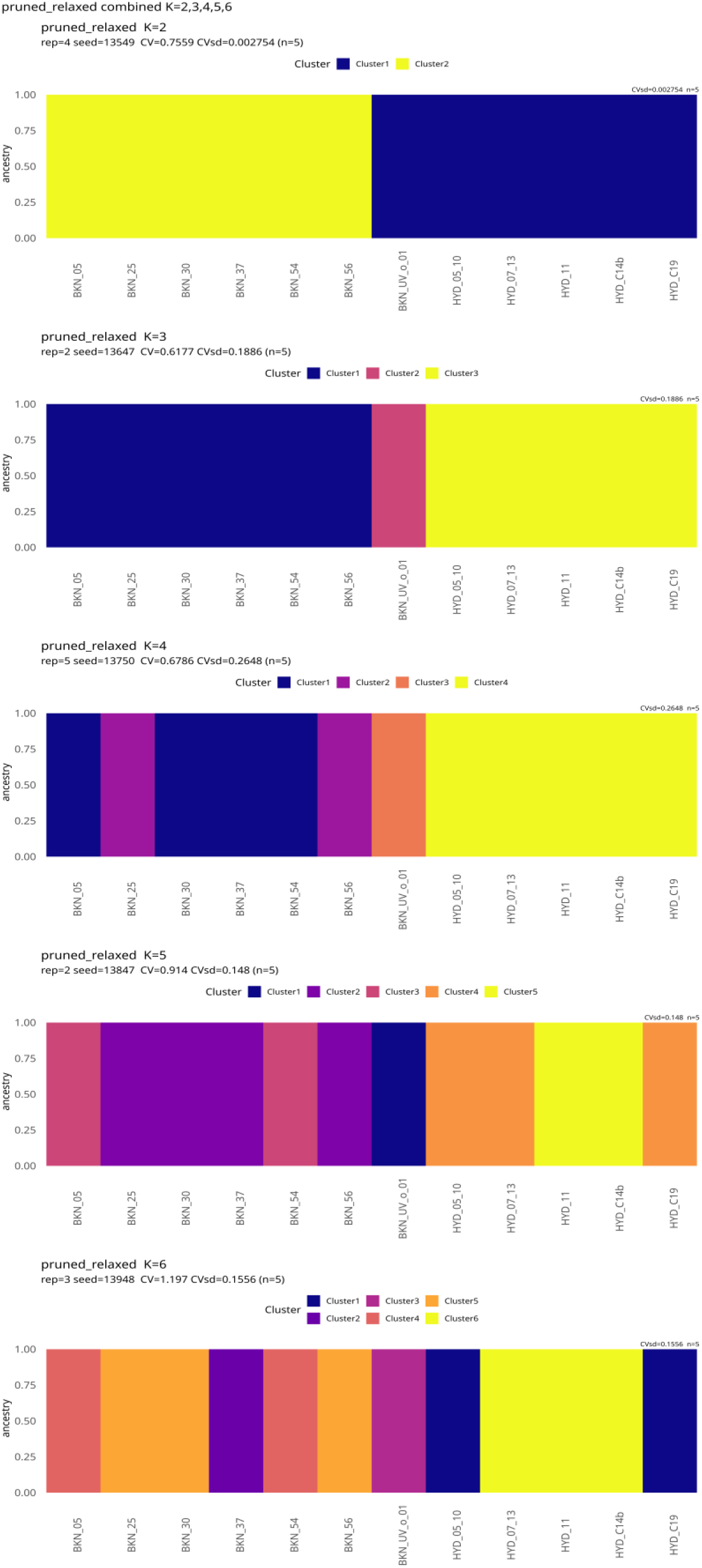
Genetic ancestry estimation using ADMIXTURE. The figure visualizes ADMIXTURE results for **K = 2-6**. Individual ancestry proportions from LD-pruned SNPs shown for K = 2-6 (replicates run per K; cross-validation used to assess stability). (ADMIXTURE settings reported in *Methods*).

**Fig S14:**
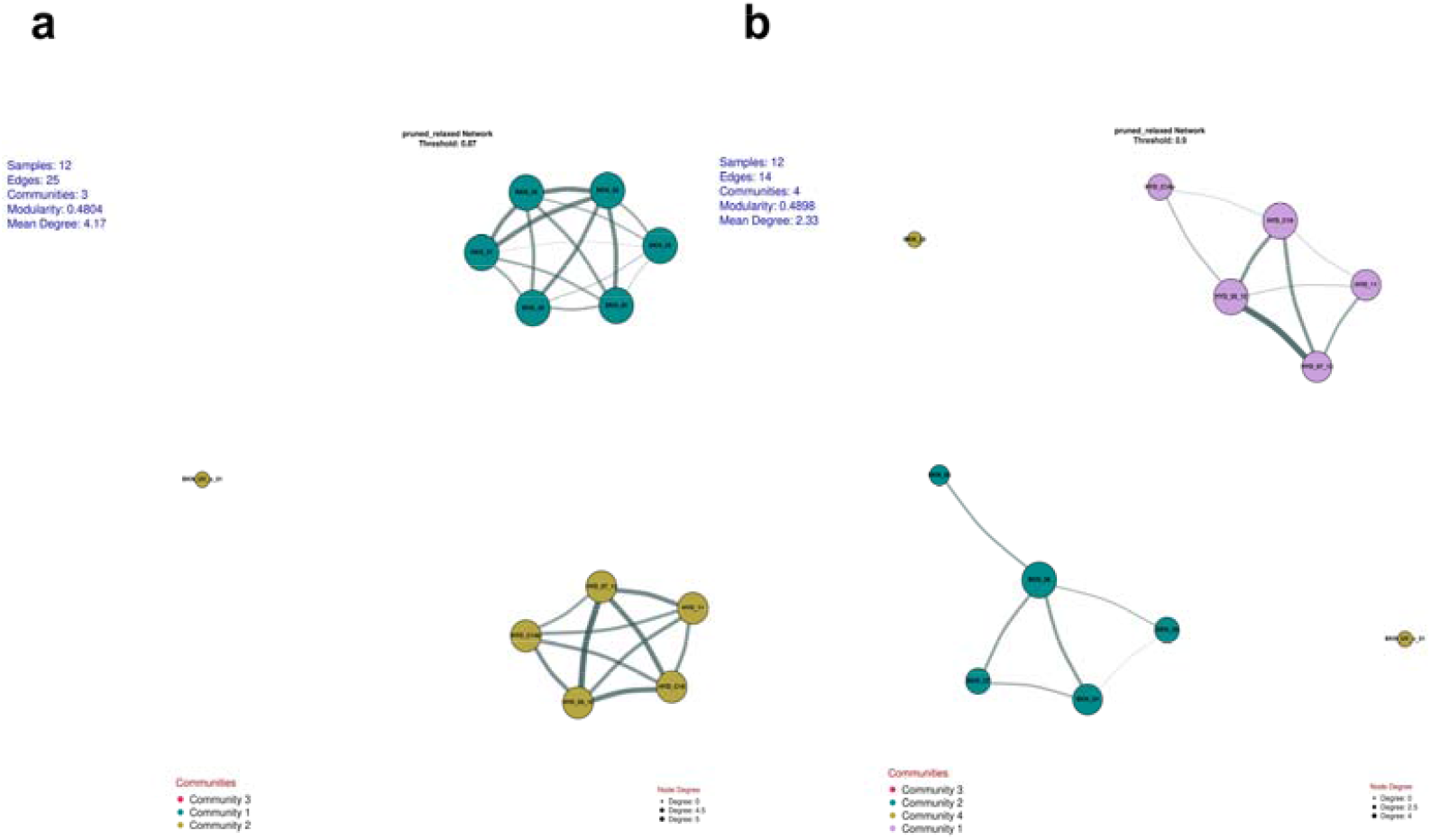
Genetic similarity networks at higher thresholds (t=0.87 and t=0.90). The behaviour of sample BKN_UV_o_01 (movement between communities depending on threshold) retraces PCA/ADMIXTURE results and suggests partial affinity to the HYD cluster or distinct ancestry.

**Fig S15:**
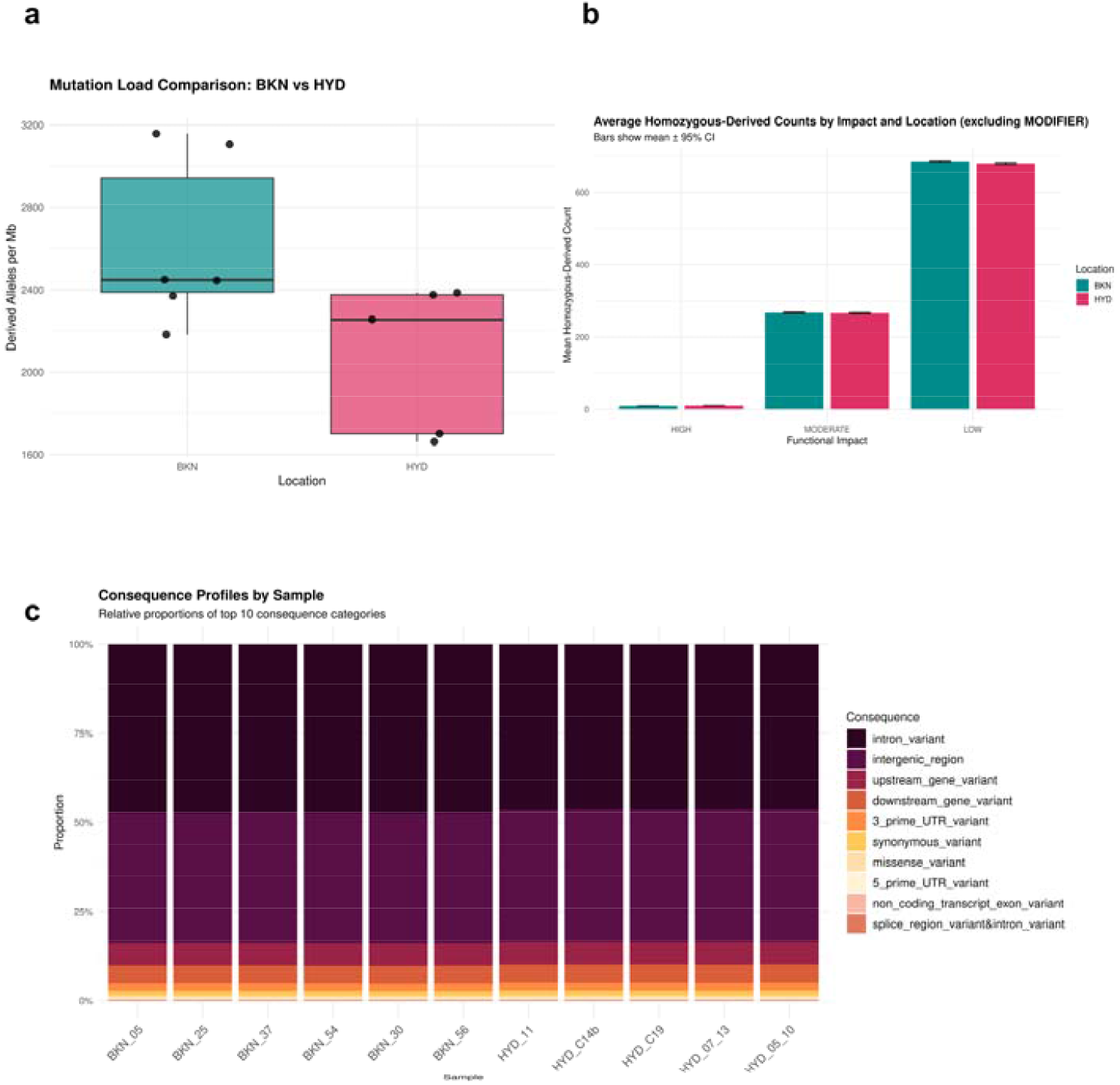
Functional composition of homozygous-derived alleles and consequence profiles. **(a) Relative proportions of homozygous-derived alleles** by impact class (LOW, MODERATE, HIGH) excluding MODIFIER (non-coding) sites — LOW and MODERATE categories comprise the bulk of coding impacts. **(b) Mean homozygous-derived counts by impact** (bars ± 95% CI) excluding MODIFIER. Together these panels reflect that the modest excess of homozygous-derived load in BKN is majorly driven by non-coding variation. **(c) Stacked consequence-profile** showing the top annotated consequence classes per sample (e.g., intronic/intergenic dominate; missense and stop_gained are rare).

**Fig S16:**
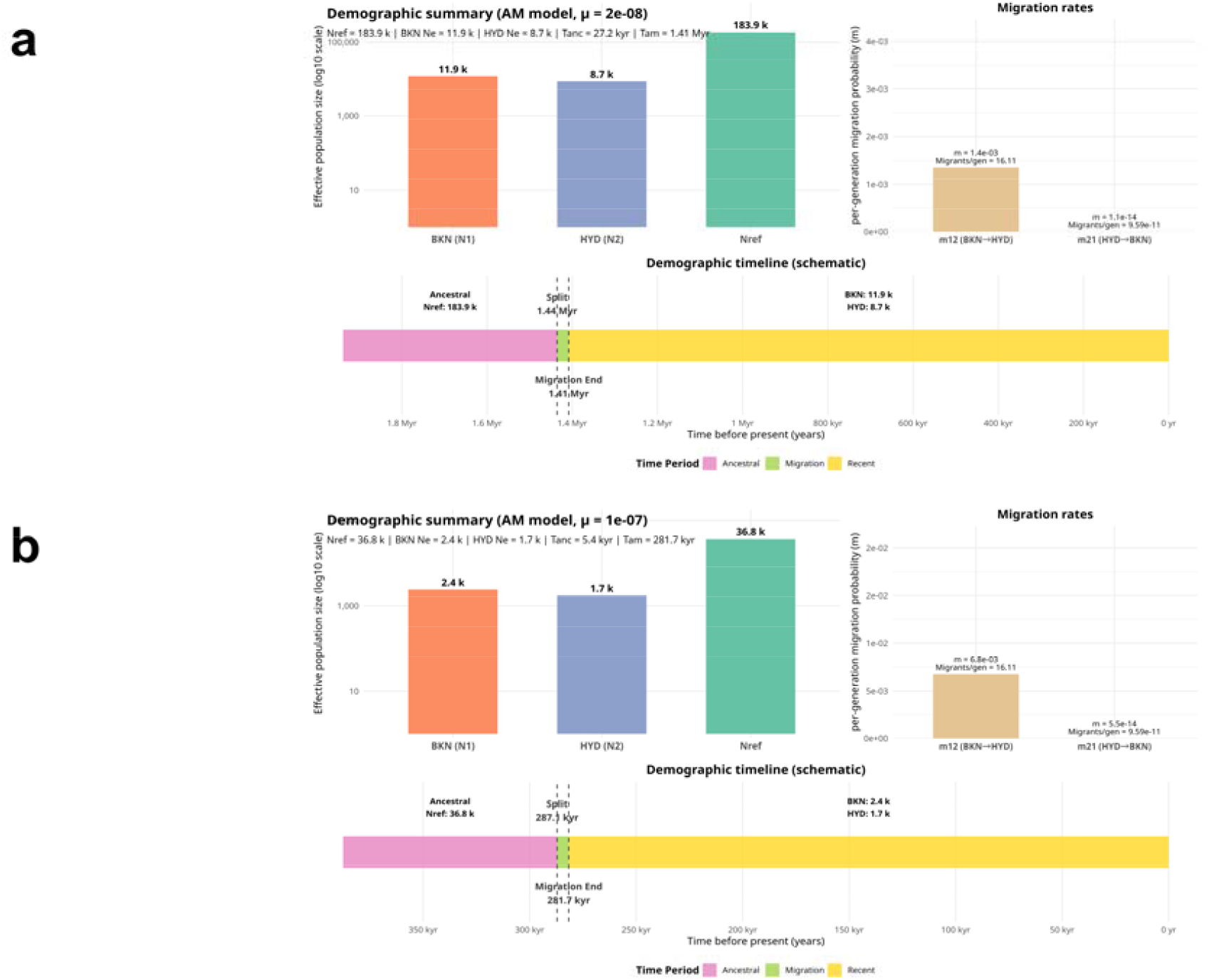
Effect of mutation rates toward inferred demographic timeline under AM model. Projections for *N*_*Ref*_ and *N*_*e*_ change for less when a higher mutation rate is used in the calculations. Panel **(a)** sketches the scenario where mutation rate is doubled from the standard assumptions while **(b)** exhibits even more radical drops in the estimated population sizes indicating the sensitivity of demographic inferences to mutation rates.

### Supplementary Tables

**Table S1:** Vulture species found in India, where two-letter codes denote the IUCN conservation status of the species: CR: Critically Endangered; EN: Endangered; NT: Not Threatened; LC: Least Concerned (LC) species. Long-term LCI and UCI stand for lower and upper limit of 95% confidence interval of modelled approximation of the change in abundance of the species in 2022-23 relative to pre-2000 values, respectively (SoIB 2023).

**Table S2**: General DNA extraction workflow summary. Note that the summary is only shown for clot-based extraction.

**Table S3**: List of samples selected based on yield and quality of genomic DNA extraction. The quantitative readings provided here were recorded from a Nanodrop spectrophotometer. (Abbreviations: BKN - Bikaner; HYD - Hyderabad).

**Table S4:** Results of the BLAST search performed to identify aligning regions of the RAG cDNA sequence in different species.

**Table S5**: RAG201-based primer pairs used for gDNA validation.

**Table S6**: Raw Data Quantification and Genotyping QC.

**Table S7**: ROH results breakdown: The table summarizes per-sample ROH tract statistics for intermediate (1000-2000 kb), old (2000-5000 kb), and ancient (>5000 kb) size categories.

**Table S8**: ADMIXTURE run results for both the stringent and relaxed LD-pruned SNP sets for K values ranging from 2-6.

**Table S9**: SNP-eff based polarized mutation load quantification results showing bootstrap summary.

**Table S10**: Group-wise mutation load estimation summary.

**Table S11**: Model performance assessed taking into account the log-likelihood for each run the and number of parameters (required to compute AIC).

**Table S12:** Demographic model-runs and parameter estimates.

**Table S13**: Bootstrap summary for demographic model-runs.

### Supplementary Methods

#### Sampling

An inclination for freshly-shed contour and tail feathers was adopted for the sampling process, and damaged, dirty, hollow or otherwise contaminated feathers were shirked (***Supplementary Table S2***). Each feather was placed in a dry paper envelope, allowed to air dry, and immediately sealed to avoid cross-contamination. The feathers were stored in the dark at ambient laboratory temperature until the date of DNA extraction (Presti et al. 2013; Ramón-Laca et al. 2018; Volo et al. 2008), which was followed up in an *in-house* facility.

#### gDNA extraction, QC and Genotyping

To validate the extracted DNA, we identified a housekeeping gene reflecting little sequence divergence within the clade, and based the primer design on sequences maximally conserved across multiple birds. The cDNA sequence of the RAG-201 gene from chicken was extracted (NCBI accession: NC_052536.1), and an NCBI-BLAST search was performed to obtain homologous sequences in six different bird species (*Pavo cristatus:* CM099179.1; *Cathartes aura:* JMFT01125012.1; *Aegypius monachus:* JANUXQ010000034.1; *Neophron percnopterus:* JBJGJM010177978.1; *Gypaetus barbatus:* CM051539.1; *Passer domesticus:* NC_087479.1) (refer to ***Supplementary Table S4***). Subsequently, an MSA was generated using Clustal v1.2.4, and primer pairs were designed (based only on the totally conserved regions) using NCBI Primer-BLAST (***Supplementary Table S5***).

Polymerase Chain Reactions (PCR) for 10 µL setups were achieved using 2X Taq DNA PCR Master Mix (*Barcode Biosciences*, HSN: *29349900*) as per the manufacturer’s instructions. The general thermal cycling program settings for 30 cycles were: denaturation at 94 °C (45 *sec*); annealing at 53.8 °C (40 *sec*); and extension at 72 °C (90 *sec*). The initial denaturation was performed at 94 °C for a minute, while the final extension was at 72 °C for 8 *mins*.

#### Dataset Curation and Workflow Finalisation

An iterative, reproducible pipeline was used to convert raw ddRAD-seq data into two analysis-ready SNP subsets: a) a non-LD pruned (full) set for diversity, heterozygosity and ROH (Runs of Homozygosity) estimates, and b) an LD-pruned set for structure and demographic inferences. The pipeline incorporated read-level quality control (QC), alignment, per-sample alignment (in BAM format) processing, joint genotyping (using GATK – haplotype caller; other variant calling pipelines were also investigated), conservative filtering, and some amount of manual curation driven by pilot-run inferences from an initial 15-sample ddRAD-seq data analysis workflow (Andrews et al. 2016; Díaz-Arce and Rodríguez-Ezpeleta 2019; Luca et al. 2011; Mastretta-Yanes et al. 2015) (for details, refer to ***Supplementary Fig. S4***).

The initial pipeline run with the complete 15-sample ddRAD-seq dataset was utilised to probe missingness, sample clustering and duplicate/kinship structure. The pilot run identified and led to the following dataset decisions that were assimilated in the final pipeline based on high-missingness sample removal, outlier (singleton) sample detection, and duplicate sample removal.

The revised pipeline (***Fig. 1***) following the recommendations from the pilot run included trim reads (fastp) alignment against the golden Eagle (*Aquila chrysaetos*) reference genome using BWA-MEM, and subsequent BAM files were coordinate-sorted and indexed. Picard MarkDuplicates was used to flag potential PCR duplicates and to generate duplicate metrics for library assessment. Because ddRAD data can yield many coordinate-identical fragments that are biologically informative (restriction-site anchored fragments), we ran GATK HaplotypeCaller with the ‘unmark duplicate’ option — a choice supported by our pilot comparison in which retaining duplicate-tagged reads increased SNP recovery (≈86 K SNPs with “unmark dup” vs ≈48 K SNPs when duplicates were strictly filtered) and reduced artefactual loss of heterozygous calls (Auwera 2013; Díaz-Arce and Rodríguez-Ezpeleta 2019; Unmark duplicate-GATK 2023). This also improved the results for kinship and heterozygosity estimates, which had earlier produced extreme values not akin to a biological context.

Joint genotyping using *GATK’s genotypeGVCFs* was followed by conservative site- and genotype-level filtering for the remaining sample dataset. The final filtered data (in VCF format) were converted to binary file format (PLINK), and three analysis sets were generated: a non-LD pruned SNP set, and two LD-pruned sets (one relaxed and one stringent using PLINK sliding-window pruning) for population structure and demography-based inferences. The option *--indep-pairwise* was used to generate the pruned sets with the following parameters for window size (number of SNPs), step size, and *r*^*2*^: stringent (100 10 0.3) and relaxed (50 5 0.4).

#### Heterozygosity and Inbreeding Assessment

PLINK’s *het* function computes heterozygosity as the proportion of heterozygous genotypes per individual relative to total genotypes, while adjusting for allele frequency estimates derived from the same dataset earlier. The inbreeding coefficient (F) was estimated as –

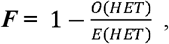

where O(HET) and E(HET) are observed and expected heterozygous genotypes, respectively.

Further, the proportion of the autosomal genome contained within RoH was calculated to estimate genomic inbreeding –

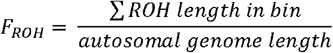

#### Population Differentiation Analysis (Pairwise FST Measure)

Pairwise FST was computed as-

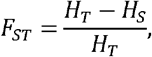

where *H*_*T*_ = total expected heterozygosity (across both populations) and *H*_*S*_ = mean expected heterozygosity within subpopulations.

#### Nucleotide Diversity

For two sequences *i* ∧ *j*, a *s* site is given by the allele counts *n*_1_,*n*_2_,…_*n*_*k*__, where *n* = ∑_*i n*_i__ is the total number of alleles observed the two estimates of genetic diversity computed are given by *π*_*site*_ is then computed as-

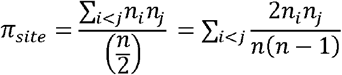

The vcftools command was used to evaluate both windowed π and per-site π for each population:

vcftools

> --gzvcf filtered_snps.vcf.gz
>
> --keep <pop_samples.keep>
>
> --window-pi <WINDOW_BP>
>
> --window-pi-step <STEP_BP>
>
> --out pop_pi

vcftools

> --gzvcf filtered_snps.vcf.gz
>
> --keep <pop_samples.keep>
>
> --site-pi --out pop_pi

π_*window*_ is estimated as-

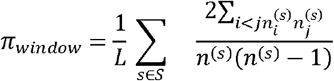

where L = length of the callable genome; S = set of polymorphic sites considered in the window; 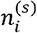 and 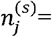 counts of alleles *i and j* for a site *s*, respectively, while *n* ^(*s*)^ denotes the number of observed alleles at site *s*.

#### Tajima’s D Estimation

Tajima’s D evaluates the difference between two measures of genetic diversity: one from the number of polymorphic loci (S) and the other from the mean number of pairwise differences (π), which are expected to be at equilibrium for a neutrally evolving population of a fixed size (Tajima 1989). It is given by-

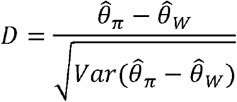

Where 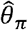 (*or just π*) is equal to 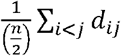 and 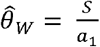 (Watterson’s diversity estimate) where *S* stands for the number of segregating sites, *a*_1_ is the scaling factor further described as 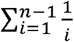 and *d*_*ij*_ indicates the number of nucleotide differences between sequences *i and j*. To obtain per-site statistics for both the 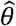 estimates, they can be divided by *L* (the length of callable genome).

#### Mutation Load Quantification

To address the biases introduced when comparing variants across species, we polarised alleles by mapping an outgroup assembly (Peregrine falcon; FASTA) to the focal reference and called the outgroup base at each focal SNP position. We implemented a three-step pipeline as reproducible shell + Python scripts for mapping and persite outgroup calling: (1) outgroup → reference mapping with minimap2 v2.30-r1287 (assembly preset -x asm5), (2) generation of a one-sample outgroup VCF by extracting reference-centred windows and parsing SAM alignment into per-site OG_BASE annotations, and (3) polarisation and mutation-load quantification using a Python script that reads the focal VCF, the outgroup calls VCF, SnpEff annotations, and a per-sample callable-bases map (extracted using the BAM files). Mapping and per-site calling also used samtools/bcftools utilities; exact commands and wrappers are archived with the pipeline.

To further reduce bias due to species composition at geographical sites and private fixed differences, we removed sites from downstream summaries if the outgroup call was missing, ambiguous (heterozygous), or below depth-thresholds; fixed interspecific differences (sites where OG_BASE consistently differs from the reference but the site is monomorphic in the focal samples) and non-segregating sites were also excluded so only confidently-polarized sites contribute to load estimates. Samples were grouped as BKN and HYD (BKN_o_UV_01 was excluded from group-level summaries). Per-sample derived-allele dosage was computed relative to the outgroup as 0 (homozygous reference), 1 (heterozygous), or 2 (homozygous). We present two complementary per-individual metrics: (i) recessive load — the sum of homozygous derived sites stratified by impact class, and (ii) additive/burden — the sum of derived-allele dosages weighted by impact (sensitivity analyses applied configurable weights, e.g., HIGH = 1.0, MODERATE = 0.5, LOW = 0.1, MODIFIER = 0.0). Each per-sample count was normalised by per-sample callable base-pairs (callable_bp) and expressed per Mb.

### Supplementary Results

#### Genomic DNA (gDNA) Extraction

The mean DNA concentration values varied greatly based on the protocol and the biological source material. The extractions following the PCI protocol yielded more consistent results for calamus tips (14.98 ± 5.68 ng/µL; n = 23), but they were nonetheless scant for genotyping and other downstream sequencing applications (***Supplementary Fig. S5***). In contrast, the PCI protocol for clot samples did not produce quantifiable DNA.

The silica kit-based extraction of calamus tip (including inferior umbilicus) also invariably failed to yield amplifiable DNA with a mean concentration of < 5 ng/µL. While the success of clot-based extractions from kits varied greatly (31.12 ± 22.98 ng/µL; n = 28), we found significant improvement when only rectrices (tail feathers) were considered (42.96 ± 20.40 ng/µL; n = 18) across all vulture species.

To confirm the avian origin of the extracted DNA, PCR amplification of one of the conserved genes of birds, namely RAG2 (Recombinase Activating gene), was carried out; successful amplification was visualized with agarose gel electrophoresis (***Supplementary Fig. S6)***. General quality check of (non-PCR amplified) extracted DNA showed, in general, poor DNA integrity of 15 of them (largely degradation), and one of them failing the QC (see ***Supplementary Fig. S7)***. The 15 samples were genotyped using the ddRADseq protocol for genome-wide SNP generation, and DNA fragment analysis during library preparation confirmed the presence of short, degraded fragments—consistent with the initial gel electrophoresis results, thus library construction was performed under high-risk conditions.

#### Pilot-run-based Curation Addressed the Grouping or Genotyping Issues

Based on the pilot run following issues were observed which were appropriately addressed: (i) **High-missingness**:-Sample BKN_24 exhibited extremely high per-sample missingness and was excluded from all downstream analyses (see the top panel in ***Supplementary Fig. S8***), (ii) **Outlier**:-Sample BKN_35 behaved as an outlier: in the structure-based analyses (see results of PCA, Genetic Networks, FST estimates and genetic ancestry in later sections), wherein it maintained a singleton grouping distinct from other BKN samples and was markedly closer to HYD samples in some metrics (can be inferred from the population divergence results discussed later). BKN_35 subsequently was renamed BKN_UV_o_01, and the reconstituted BKN outlier group BKN_o was excluded from group-level statistics (it was, however, retained in individual-level summaries). (iii) **Duplication**:-Kinship and relatedness analysis revealed two biologically duplicate pairs a) HYD_05 & HYD_10 and b) HYD_07 & HYD_13 (see ***Supplementary Fig. S9***). Rather than discarding the duplicates, the raw sequence files for each duplicate pair (at reads level) were merged, and treated as a single sample for variant calling, aimed at ameliorating effective coverage and have already been shown to improve variant discovery in duplicated/exomic replicates (refer to ***Supplementary Fig. S8*** for genotype-level comparison for before and after data curation) (Zhang et al. 2014).

#### ROH Size Spectrum Reveals Overall Higher Recent Autozygosity in BKN Samples

While inferring genomic inbreeding, the F_ROH_ values reiterated severe long-term reductions in the effective population size and prolonged population isolation of the sampled groups — a genomic context that can raise the realised risk of inbreeding depression and compromised adaptive potential (Kardos et al. 2016; Keller and Waller 2002; Frankham et al 2010). Although captive populations are often predicted to have elevated inbreeding due to smaller breeding pools and managed founder sets (Frankham et al 2010), the small difference in the values between sites may instead reflect the different species composition (BKN includes Egyptian and Eurasian griffon vultures; HYD houses White-rumped vultures) and/or differences in connectivity and local demography; however given the modest effect size and limited sample numbers, we interpret this as suggestive rather than definitive.

Although inbreeding estimates from PLINK corroborate the findings from ROH analysis suggesting inflated inbreeding levels, they differ greatly in magnitude for methodological reasons: PLINK’s heterozygosity-based F is sensitive to SNP ascertainment, the set of loci used and to missing data; ROH based F is a direct measure of long homozygous tract and is generally more informative about autozygosity and recent inbreeding (Ceballos et al. 2018a,b; Purcell 2007). Because our ROH analysis and -*het* results are consistent qualitatively, we treat the PLINK F values here as confirming an extreme reduction of heterozygosity, but we caution against interpreting the values numerically, and instead, the ROH-based F_ROH_ should be preferred for exposition of inbreeding history and magnitude (McQuillan 2008).

#### Species Composition in Different Enclosure Settings (Open/Captive) Drive Observed Genetic Clustering

##### PCA

Samples from the BKN site cluster tightly and project to PC1 negative values, while HYD samples cluster separately (PC1 positive and PC2 slightly negative; see ***Fig. 3a***). As mentioned earlier, the outlier sample (BKN_UV_o_01) was distinct from the other two clusters along PC1 (projects closer to HYD on PC3; see ***Supplementary Fig. S12***).

##### Pairwise F_**ST**_

The groupwise per-site summary (outlier excluded) reported mean and median F_ST_ to be 0.2626 and 0.079, respectively, across 8,503 sites — indicating a skewed per-site distribution with a subset of highly differentiated loci. This could also stem from small sample sizes and/or site filtering/ascertainment that can inflate the mean of per-site F_ST_. For robust statements about genome-wide divergence, we therefore rely on the overall picture from all the population-group-based analyses (PCA, ADMIXTURE, F_ST_, and genetic similarity networks) rather than on the single raw mean value alone.

#### Negative Tajima’s D and Low π Consistent with Recent Bottlenecks

Comparisons of nucleotide diversity with typical wild avian species place the samples in the same class as critically endangered or long-bottlenecked taxa (e.g. kakapo, black robin), where π is often orders of magnitude lower than widespread wild birds (Dussex 2021; Seth 2022). Collectively, the design is consistent with recent demographic disconcertion (bottleneck and/or incomplete recovery) and/or pervasive purifying selection acting on linked sites (Li and Durbin 2011; Nielsen 2005). HYD is a captive assemblage inhabited by White-rumped vultures, while BKN holds a mixed cohort of species. Consequently, the slightly inflated mean π is likely a reflection of mixed founder history, and species composition and sampling design rather than a considerably larger N_e_: captive holdings may retain allelic forms from multiple founders or lineages, escalating observed π relative to a small wild cohort (Dussex 2021; Woodworth et al. 2002; Frankham et al 2010).

#### Reconciling negative D with high autozygosity (F_ROH_)

We observe a pattern exhibited when a population endures a recent bottleneck or severe founder event: bottlenecks increase autozygosity while generating an SFS with many rare variants. For example, if BKN experienced a severe founder event followed by a recent influx of rare mutations (or retained many rare alleles from diverse founders), windowed D may be negative even when F_ROH_ is high (Li and Durbin 2011). Although we mitigated the ascertainment and sparse coverage biases associated with ddRAD SNP sets by tuning window sizes to SNP density, residual bias remains, likely affects the SFS, and can also push Tajima’s D towards negative values. (Nielsen 2005). This technical bias is expected to influence both populations but may differ between them due to species composition or differences in per-population SNP calling rates. However, we advise against interpreting the results at face value and caution that comparisons between sites should be drawn in light of the population differentiation/divergence and ROH results because species composition and sample size differences can shift summary statistics.

The combined evidence — low π, substantial ROH-burden, negative Tajima’s D, and clear site partitioning LJ is alarming for both biological resilience and long-term adaptive potential (Hedrick and Garcia-Dorado 2016; Frankham et al 2010). In particular:

- **HYD (captive White-rumped assemblage)**: modestly higher π but similar F_ROH_ profile implies founders may have originated from multiple source lineages, yet inbreeding has accumulated; managed breeding could maintain diversity but requires active genomic monitoring (Seth 2021; Frankham et al 2010).
- **BKN (open enclosure with Egyptian + Griffon vultures)**: more negative D and higher F_ROH_ signify stronger recent bottleneck or constrained effective gene flow among the resident species, and the presence of long ROH tracts heightens the susceptibility to risks linked to inbreeding depression (Hedrick and Garcia-Dorado 2016; Lavanchy and Goudet 2023; Robinson 2019).

#### Demographic Reconstruction Reveals Early Divergence and Prolonged Population Isolation

The post-split population contraction explains the strong genome-wide signatures we note independently: both populations carry elevated runs of homozygosity and high inbreeding, with BKN consistently exhibiting more extreme levels and a higher predicted burden of deleterious variation. Shrunken N_e_ after divergence expounds the genomic signatures of increased genetic drift and reduced efficacy of purifying selection, reflected in the observed accumulation of mildly deleterious alleles in BKN.

Additionally, the fitted migration parameters are highly asymmetric. Converted into migrants per generation applying the standard ∂a∂i scaling (and the inferred **θ**), we find the scaled m12 to be large, while m21 to be essentially zero. Interpreted in demographic terms, this implies a historical pulse of appreciable gene flow into the BKN lineage during the migration period, followed by subsequent isolation. Such an asymmetric pattern of directional exchange during the post-split migration window entails historical connectivity biased toward BKN as a sink and HYD as a source. This can also produce allele sharing despite later reproductive isolation and may explain some patterns of allele frequency similarity alongside localised inbreeding in BKN. Note, however, the extremity in scaled migration parameters can partly reflect model parametrisation and the confines of folded SFS; we therefore emphasise the qualitative asymmetry rather than absolute numeric magnitude.

